# Characterization of striatal dopamine projections across striatal subregions in behavioral flexibility

**DOI:** 10.1101/2021.09.18.460922

**Authors:** R.K. van der Merwe, J.A. Nadel, D. Copes-Finke, S. Pawelko, J.S. Scott, M Fox, C. Morehouse, M. Ghanem, R. McLaughlin, C. Maddox, G. Malaki, A. Turocy, X. Jin, C.D. Howard

## Abstract

Behavioral flexibility is key to survival in a dynamic environment. While flexible, goal-directed behaviors are initially dependent on dorsomedial striatum, they become dependent on lateral striatum with extended training as behaviors become inflexible. Similarly, dopamine release shifts from ventromedial to lateral striatum across learning, and impairment of lateral dopamine release disrupts habitual, inflexible responding. This raises the possibility that lateral dopamine release is a causative mechanism in establishing inflexible behaviors late in training, though this has not been directly tested. Here, we utilized optogenetics to activate dopamine terminals in dorsal medial (DMS), dorsal lateral (DLS), and ventral (NAc) striatum in DATcre mice to determine how specific dopamine subpopulations impact behavioral flexibility. Mice performed a reversal task in which they self-stimulated DMS, DLS, or NAc dopamine terminals by pressing one of two levers before action-outcome lever contingencies were reversed. Consistent with presumed ventromedial/lateral striatal function, we found that mice self-stimulating ventromedial dopamine terminals rapidly reversed lever preference following contingency reversal, while mice self-stimulating dopamine terminals in DLS showed impaired reversal learning. These impairments were characterized by more regressive errors and reliance on lose-stay strategies following reversal, suggesting reward insensitivity and overreliance on previously learned actions. This study supports a model of striatal function in which dorsomedial dopamine facilitates goal-directed responding, and dorsolateral dopamine release is a key mechanism in supporting the transition toward inflexible behaviors.

## Introduction

The ability to rapidly update previously learned action-outcome associations is a key feature of flexible, goal-directed behavior. Reversal learning, or switching between competing and often opposing reward-related behaviors, is one commonly studied form of cognitive flexibility (Izquierdo et al., 2017). As an example of reversal learning, international travel can often require a driver to rapidly update on which side of the road they are driving, and it can require great cognitive effort to not default to previously learned and deeply-ingrained behaviors (i.e., driving on the more familiar side). Accordingly, several diseases associated with reduced cognitive flexibility, including obsessive-compulsive disorder, drug addiction, Parkinson’s disease, and schizophrenia, are characterized by impairments in reversal learning (Clark et al., 2004; Gillan et al., 2011; Leeson et al., 2009).

The striatum, the primary input nucleus of the basal ganglia, is thought to guide behavioral selection (Mink, 2003). Recent work in animal models has suggested distinct roles of dorsal striatal subregions in behavioral flexibility, with medial aspects promoting and lateral aspects impairing flexibility. Lesioning dorsomedial striatum produces deficits in goal-directed behavior (Yin et al., 2005), while lesioning dorsolateral striatum disrupts the emergence of inflexible, habitual behavior (Yin et al., 2004; Hilario et al., 2012). Activity (Gremel & Costa, 2013) and synaptic strength (O’Hare et al., 2016) in dorsomedial and dorsolateral striatum are associated with flexible and habitual responding, respectively. Similarly, lesions of dorsomedial striatum impair reversal learning (Castañé et al., 2010), while optically silencing lateral striatum enhances reversal learning (Bergstrom et al., 2018), suggesting that lateral striatum establishes inflexibility by promoting well-ingrained behavioral patterns after repeated exposure to action-outcome associations. In contrast to dorsal striatum, ventral striatum, and specifically nucleus accumbens core (NAc), is necessary for acquisition of new skills and may be implicated in behavioral flexibility. Lesions of the NAc core disrupt aspects of spatial reversal (Stern & Passingham, 1995), though other studies have reported more modest effects of NAc core lesions on flexibility (Floresco et al., 2006). Based on these findings, a model has emerged positing that flexible behavior reflects dependence on ventro-medial striatal circuits, which shifts to lateral striatum as tasks become well learned and inflexible (Thorn et al., 2010; Yin et al., 2009).

Striatum receives dopamine inputs from midbrain, which are required for behavioral flexibility during reversal of action-outcome contingencies. Dopamine is a key neurotransmitter in the learning of associations and their predictive cues (Schultz, 1998; Tsai et al., 2009). Impairments in dopamine transmission slow behavioral reversal (Clarke et al., 2011; Radke et al., 2019) and dopamine D2 receptor availability is associated with rate of reversal learning in monkeys (Groman et al., 2011). Phasic increases in dopamine concentration provide positive feedback about outcomes, and predict an animal’s ability to reverse (Klanker et al., 2015). Accordingly, optogenetic activation (Adamantidis et al., 2011) or photosilencing (Radke et al., 2019) of dopamine neurons facilitates or impairs reversal, respectively. Consistent with the functional divide between dorsomedial and dorsolateral striatum, lesioning dopamine inputs to DMS specifically impairs reversal (Grospe et al., 2018), while lesioning dopamine inputs to DLS impairs inflexible responding and renders animals goal-directed (Faure et al., 2005; Faure et al., 2010). Interestingly, the pattern of dopamine release across subregions is dynamic, with dopamine release in ventromedial striatum occurring early in learning, and dopamine release in lateral striatum emerging late in learning as behaviors become well-learned (Willuhn et al., 2012; Radke et al., 2019). Further, dorsomedial and dorsolateral dopamine neurons are composed of unique populations that carry distinct information (Lerner et al., 2015). Taken together, this raises the possibility that the onset of dopamine release in dorsolateral striatum is a key mechanism in the transition from flexible to inflexible responding across learning, though this possibility has not been directly tested.

To test this hypothesis, we implanted fiber optics into DMS, DLS, or NAc core of mice that express channelrhodopsin-2 in dopamine neurons. Mice performed an intracranial self-stimulation (ICSS) reversal task, in which they received optogenetic stimulation at implantation site upon pressing one of two levers that was initially active. After demonstrating preference for the initially active lever for a period of five days, the active lever was switched and mice had to reverse lever preference to the second lever in order to receive laser stimulation. Mice preferred the active lever regardless of stimulation site, but only mice with fiber optics targeting dorsomedial and ventral striatum were able to fully reverse their preference in favor of the secondarily active lever following contingency reversal. Consistent with this, mice with fiber optics targeting dorsolateral striatum made more errors that tended to be regressive, rather than perseverative, and they relied more heavily on lose-stay strategies following reversal, suggesting self-stimulation of DLS biases mice toward previously reinforced actions and reduces sensitivity to more proximal reinforcing outcomes. Taken together, these results suggest that behavioral flexibility is mediated by striatal subregion-specific dopamine activity, with ventral and dorsomedial dopamine promoting flexible responding, and dorsolateral dopamine promoting inflexibility.

## Materials and methods

### Animals

All experiments were approved by the Oberlin College Institutional Animal Care and Use Committee and were conducted in accordance with the National Institutes of Health’s Guide for the Care and Use of Laboratory Animals and Animal Research: Reporting of In Vivo Experiments (ARRIVE) guidelines. Mice were maintained on a 12 hr/12 hr light/dark cycle and were provided *ad libitum* access to water and food. Experiments were carried out during the light cycle using a total of 36 DAT^IRES*cre*^ (JAX: 006660) and DAT^IRES*cre*^ X Ai32 *Rosa*^*ChR2(H134R)-EYFP*^ (JAX: 024109) mice ranging from 2 to 6 months of age. DATcre mice preferentially express Cre in dopamine neurons (Bäckman et al., 2006), such that DATcre mice crossed with Ai32 mice express Channelrhodopsin-2 (ChR2) in dopamine neurons (Madisen et al., 2012; B. O’Neill et al., 2017; Howard et al., 2017; Figure 1A+C).

**Figure 1.**
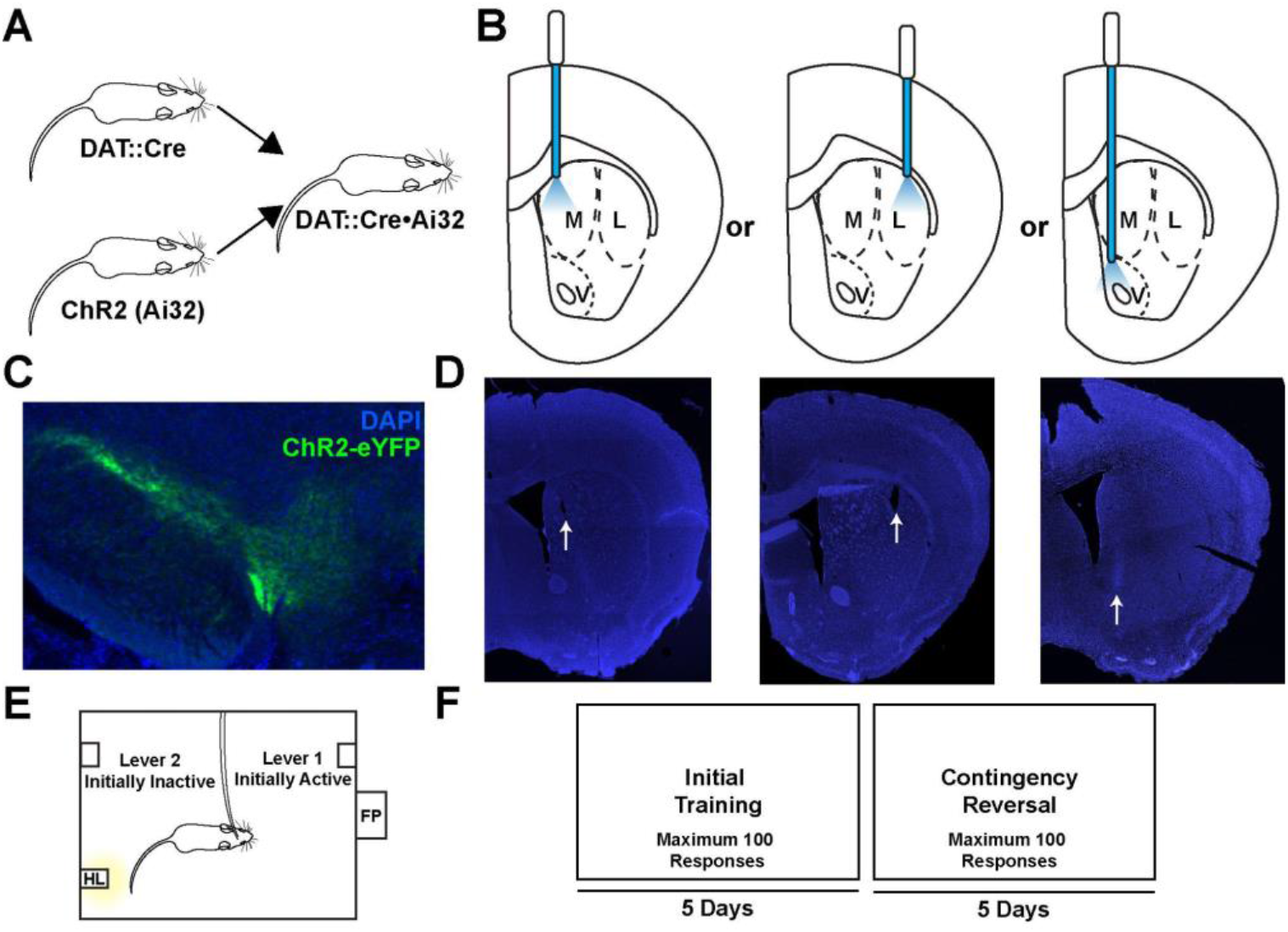
Subjects and experimental design. **A**. Breeding scheme. DATcre^+/-^ mice were crossed with Ai32^+/+^ mice to establish DATcre^+/-^ x Ai32^+/-^ mice. **B**. Mice were chronically implanted with fiber optics targeting dorsomedial (M, left), dorsolateral (L, center), or ventral striatum (V, nucleus accumbens core; right). **C**. DAT x Ai32 mice express ChR2 selectively in SNc and VTA. **D**. Representative fiber optic implantation sites for DMS (left), DLS (center), and NAc (right). **E**. Operant chamber configuration for Ai32 experiments. Two levers were extended on either side of the operant chamber. **F**. Schematic of experimental design. Mice could press either lever a total of 100 times each day, and following five days of meeting criteria for preferring the active lever (see Methods), the contingency was reversed, and the active lever became inactive and *vice versa*, and mice were tested for an additional five days.

### Reagents

Isoflurane anesthesia was obtained from Patterson Veterinary (Greeley, CO, USA). Sterile and filtered phosphate buffered saline (PBS, 1X) was obtained from GE Life Sciences (Pittsburgh, PA, USA). Unless otherwise noted, all other reagents were obtained through VWR (Radnor, PA, USA).

### Stereotactic Surgery and Viral Injections

Mice were anaesthetized with isoflurane (4% at 2 L/sec O2 for induction, 0.5-1.5% at 0.5 L/sec O2 afterward) and then placed in a stereotactic frame (David Kopf Instruments, Tajunga, CA, USA). The scalp was shaved and sterilized with povidone iodine and an incision was made in the scalp. The skull was scored with Optibond (Patterson Dental) and holes were drilled above the striatum or midbrain dopamine neurons. DATcre mice received 1µl injections of virus encoding ChR2 (AAV5-EF1-DIO-hChR2(H134R)-EYFP-WPRE-pA, UNC Viral Vector Core) targeting substantia nigra pars compacta (SNc; -3.2 AP, ±1.5 ML, -3.8 DV). A 5µL syringe needle (Hamilton) was lowered to the DV coordinate over 2 minutes and held in place for 1 min before the start of injection. Injections took place over 5 min, the needle was left undisturbed in the brain for 5 min after the completion of virus delivery, and the needle was then removed over the course of 5 minutes. Both virally-injected DATcre and DATcre x Ai32 mice received bilateral fiber optic implants to allow optogenetic activation of dopamine terminals. Virally-injected DATcre received fiber optics targeting dorsomedial striatum (DMS; +0.8 AP, ±1.0 ML, -2.3 DV) or dorsolateral striatum (DLS; +0.8 AP, ±2.25 ML, -2.3 DV) immediately following injection. DATcre x Ai32 mice received fiber optics in DMS or DLS at the same location listed above, or in the nucleus accumbens (NAc; +1.2 AP, ±1.1 ML, -4.0 DV), but received no viral injection. Fiber optic implants were secured to the skull with a skull screw and dental cement (Patterson Dental). Mice were allowed to recover for a minimum of 2 weeks (Ai32 mice) or 3 weeks (virally-injected mice) before behavioral experiments took place.

### Fiber Optic Implants

Fiber optic implants were custom fabricated and were comprised of 0.39 NA, 200 µm core Multimode Optical Fiber (ThorLabs) inserted into a multimode ceramic zirconia ferrules (1.25mm OD, 230um ID; SENKO). The fiber optic was affixed in the ferrule with two-part epoxy (353ND; Precision Fiber Products). Each end of the fiber optic was polished using fiber optic sandpaper (ThorLabs) and functionality was tested ensuring minimal loss of light power and even output prior to implantation.

### Intracranial Self-Stimulation Reversal Task

Behavioral sessions took place in a Med Associates behavioral chamber housed inside a sound-attenuating cubicle. The behavioral chamber was outfitted with two levers, a house light, and a food port. Two distinct orientations of levers were utilized in these experiments. For virally-injected DATcre mice, levers were oriented on either side of the food port (Supplemental Figure S1E). For subsequent DATcre x Ai32 experiments, the lever closest to the door was later moved to the back of the opposite wall to make proximity to the door consistent across both levers (Figure 1E). At the beginning of behavioral sessions, mice were briefly anesthetized with isoflurane (4%, 2 l/min O2) and fiber optic implants were connected to fiber optic leads inside the behavioral chamber, which were connected to a diode-pumped single-state laser (Laserglow, 473nm) by a commutator (Doric Lenses) to allow for free rotation. Mice were placed on a ≥10 day optogenetic intracranial self-stimulation (ICSS) task (Figure 1F and Supplemental Figure S1F for Ai32 and virally-injected mice, respectively). Here, all sessions began with the illumination of the house light and extension of both levers. Depressing one of the two levers resulted in laser onset while the opposite lever had no effect (see **Laser Stimulation** for stimulation details). Mice were trained to self-stimulate for at least 5 days with no changes in contingency. During this five day period, mice were required to meet three criteria: 1. A minimum of 10 presses on the active lever, 2. Preference (>50% of all presses) for the active lever on at least 4 of the 5 days, and 3. Preference for the active lever on the fifth day of training. Mice that failed to reach these criteria were provided up to 10 additional training sessions until the criteria was achieved, and, if they failed to reach this criteria, were excluded from the study. To test response flexibility between self-stimulation contingencies, mice that met these criteria underwent reversal on day six, in which the active and inactive lever contingencies were reversed. Therefore, pressing the formerly active lever now had no effect and the formerly inactive lever triggered laser onset. Virally-injected DATcre mice were allowed unrestricted lever pressing during 90 min sessions. To partially control for variable press rates seen in virally-injected mice, DATcre x Ai32 mice were capped at a total of 100 presses on either lever, and sessions were terminated when 100 presses were made or after 60 minutes - whichever came first. At the end of each behavioral session, the house light was turned off and both levers were retracted.

### Laser Stimulation

Prior to all behavioral sessions, laser output was calibrated to 10 mW for virally-injected DATcre and 5 mW for DATcre x Ai32 mice from the end of fiber optic leads using an optical power meter (ThorLabs). During self-stimulation trials, virally-injected DATcre mice received 50Hz, 50p, 10ms pulse duration laser stimulation following each active lever press; while DATcre x Ai32 mice received 30Hz, 30p, 24ms pulse duration stimulus trains. A subset of DATcre x Ai32 mice (2 DMS, 2 DLS) received 1 sec laser pulses following each active lever press.

### Histology

After finishing all behavioral tests, mice were deeply anesthetized with isoflurane (4%, 2 l/min O2) and transcardially perfused with 0.9% saline and 4% paraformaldehyde (PFA). Brains were removed and allowed to post-fix in 4% PFA at 4°C for 24 hours. Then, brains were transferred to a 30% sucrose solution at 4°C for at least 48 hours before sectioning. Brains were sectioned on a freezing microtome into 30 μm coronal sections, and stored in cryoprotectant at 4°C. To characterize Cre expression in DATcre mice, one DATcre mouse was injected with AAV8-hSyn-DIO-mCherry (Addgene, cat#50459) in midbrain (SNc; -3.2 AP, ±1.5 ML, -3.8 DV; see **Stereotactic Surgery and Viral Injections** above). This tissue was washed 2×15 minutes in Tris buffered saline (TBS), and blocked in 3% horse serum and 0.25% Triton X-100 prior to antibody staining for 1 hour. Sections were then incubated with 1:500 diluted primary antibodies of anti-TH polyclonal rabbit antibody (Cell Signaling, cat#2792S) for 24 h at 4°C on a shaker. Following incubation, sections were washed 2×15 minutes in TBS to remove excess primary antibody, then incubated with TBS and 3% horse serum and 0.25% Triton X-100 before being incubated with Alexa Fluor© 488 Anti-Rabbit IgG (Cell Signaling, cat#4412S, diluted 1:250) for 2 hours at room temperature. Tissue was then washed 2×20 minutes in TBS to reduce background staining. Slices were subsequently floated in 0.1M phosphate buffer and mounted on slides. After drying, sections were mounted (Aqua-Poly/Mount, Polysciences, 18606-20) with DAPI (VWR, IC15757410; 1:1000). Slides were kept at RT overnight before moving to 4°C for long-term storage. Sections were imaged using a Leica DM4000B fluorescent microscope or a Zeiss LSM 880 confocal microscope.

### Data Analysis

Lever preference was defined as presses on either lever/total presses in session *100. Errors were defined as presses on the inactive lever and were counted and averaged across early reversal (days 1 and 2 following contingency reversal), and late reversal (days 3, 4, and 5 following contingency reversal) for each subject before being averaged across groups. Perseverative errors were defined as inactive lever presses following contingency reversal before the now-active lever was sampled each day. Regressive errors were defined as presses on the now inactive lever following first sampling of the now-active lever each day. Transitions were categorized as ‘lose’ if an inactive press was followed by an inactive press (lose-stay) or an active press (lose-shift), or ‘win’ if an active press was followed by an inactive press (win-shift) or active press (win-stay). Presses were calculated using Excel (Microsoft). Win/lose, shift/stay transitions were counted and verified using Matlab software MATLAB (R2018b, Mathworks). Drawings in Figure 1A,B,E,F and Supplemental Figure S1A,B,E,F were made by CDH in Adobe Illustrator CC 2019 (Adobe).

### Statistical Analysis

Statistical analysis was performed by GraphPad Prism 7.04 (GraphPad). Presses and lever preferences were compared using Two-Way Repeated Measures ANOVA with Geisser-Greenhouse corrections and *post hoc* Fisher’s LSD tests. Early vs Late reversal error analyses were compared using Two-way Repeated Measures ANOVA with *post hoc* Sidak multiple comparisons tests. For all tests, significance was defined as p ≤ 0.05 (*), and marginal significance was defined as p ≤ 0.1 (#).

## Results

### Optogenetic dopamine self-stimulation and reversal across striatal subregions

To probe the contribution of dorsal medial, lateral, and ventral dopamine projections to the striatum in behavioral flexibility, we crossed DAT^IRESCre^ mice (Bäckman et al., 2006) with Ai32 *Rosa*^*ChR2(H134R)-EYFP*^ mice (DAT x Ai32) to drive Cre-dependent expression of channelrhodopsin-2 (ChR2) in dopamine neurons (Madisen et al., 2012; O’Neill et al., 2017; Howard et al., 2017; Figure 1A+C). These mice were deeply anesthetized and implanted with fiber optics targeting either dorsomedial striatum (DMS), dorsolateral striatum (DLS), or nucleus accumbens (NAc) to activate distinct projections of dopamine inputs (Figure 1B+D, see methods). In an earlier pilot group, we employed DAT^IRESCre^ (DATcre) mice that were anesthetized and injected with AAV encoding Cre-dependent ChR2 in dopamine neurons (Supplemental Figure S1A+C). These mice were then chronically implanted with fiber optics targeting DMS or DLS (Supplemental Figure S1B+D). Data for DATcre x Ai32 mice are found in main figures throughout this text, and data for virally-injected DATcre mice are found in supplementary figures matching main figure numbers. These groups were not collapsed, as the behavioral task and operant box layout between these two groups differed (see Methods, Figure 1E-F and Supplemental Figure S1E-F).

2-3 weeks following surgery, fiber optic implants were connected to fiber optic leads capable of delivering blue (473 nm) laser light, and mice were placed in a Med Associates behavioral chamber with two retractable levers and a house light (Figure 1E; Supplemental Figure 1E). Mice performed an intracranial self-stimulation reversal schedule: they had access to two levers, and pressing one of these levers led to delivery of laser light while pressing the other lever had no effect, and after 5 days, this contingency was reversed (Figure 1F, Supplemental Figure S1F). To control for differences in rewards earned, which can modify flexibility (Adams & Dickinson, 1981), DAT x Ai32 mice were limited to 100 cumulative presses per session on either lever during the initial training and reversal phases. In contrast, virally-injected mice were allowed unrestricted pressing for both training and reversal. During the training phase, both DAT x Ai32 mice and virally-injected DATcre mice responded significantly more on the active lever relative to the inactive lever (two-way repeated-measures ANOVA; DMS, significant effect of lever, F_(1, 14)_ = 18.04, p <0.001, no significant effect of time or lever x time interaction, significant *post hoc* test for all points, p<0.05; DLS, significant effect of lever, F_(1, 12)_ = 9.839, p <0.01, no significant effect of time or time x lever interaction, significant *post hoc* tests at day 1+2, p <0.05, trending effect at days 3+5, p<0.1; NAc, significant effect of lever, F_(1, 14)_ = 9.927, p <0.01 no significant effect of time or time x lever interaction, significant *post hoc* tests at days 1+2, p <0.05, trending effect at days 3-5, p <0.1; Figure 2 A-C; Supplemental Figure S2A+B). Similarly, when normalized to the total number of lever presses, mice displayed a significant preference for the active over the inactive lever (DMS; two-way repeated-measures ANOVA, significant effect of lever, F_(1, 14)_ = 78.74, p <0.0001, and time x lever interaction, F_(4, 56)_ = 3.407, p <0.05, no significant effect of time; DLS; significant effect of lever, F_(1, 12)_ = 41.04, p <0.0001, no significant effect of time or time x lever interaction; NAc; significant effect of lever, F_(1, 14)_ = 67.4, p <0.0001, no significant effect of time or time x lever interaction; significant *post hoc* tests for all points tested across DMS, DLS and NAc, p<0.05; Figure 2 D-F; Supplemental Figure S2C+D). Taken together, terminal stimulation of dopamine neurons was reinforcing across all 3 implantation sites in both Ai32 and virally-injected DATcre mice.

**Figure 2.**
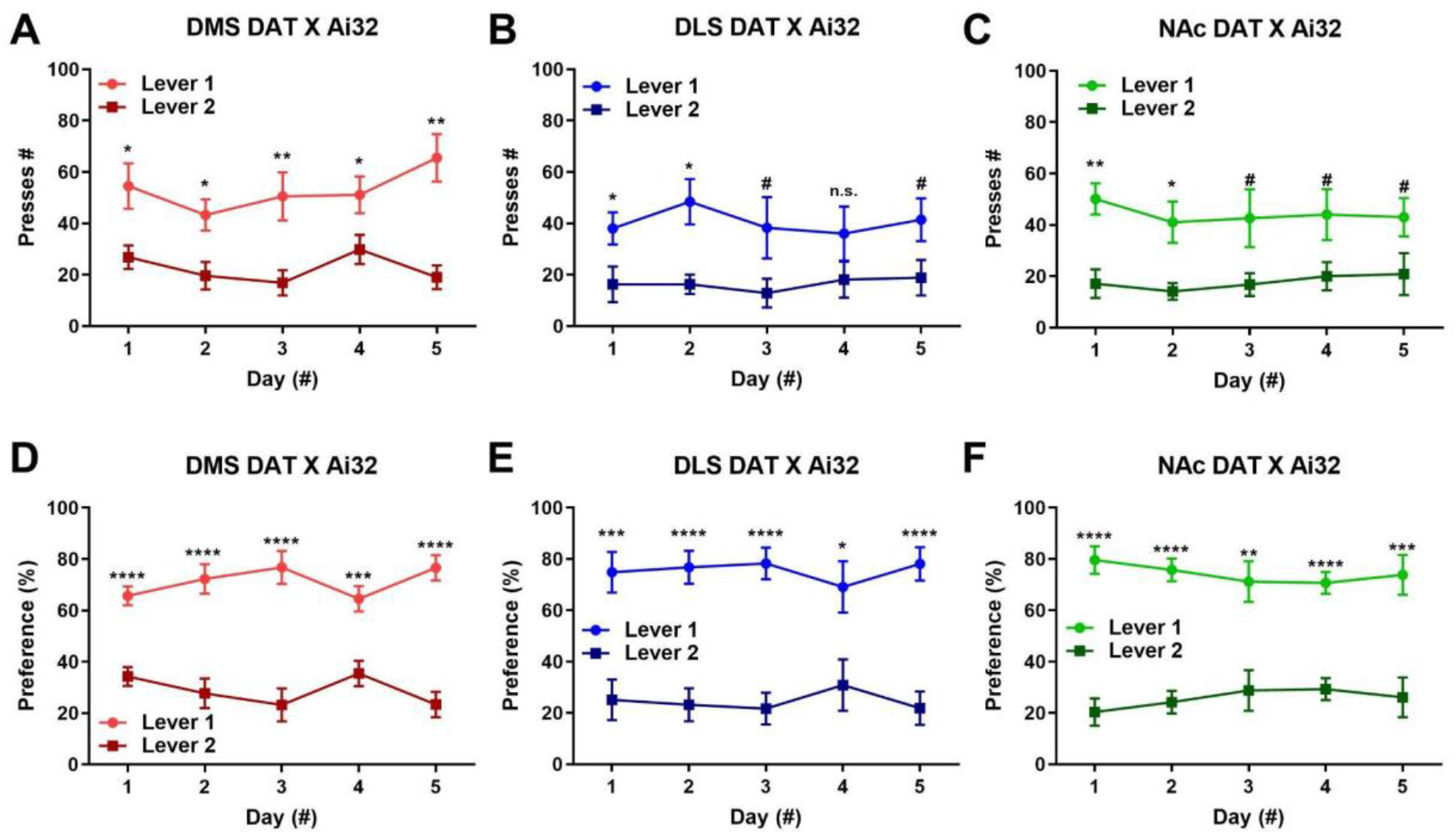
Responding during the initial training phase across implantation sites. Total press counts and preference (presses on either lever / total presses * 100) for mice self-stimulating dopamine terminals in DMS (**A+D**), DLS (**B+E**), and NAc (**C+F**) across the five training days. Mean ± SEM, ^#^p>0.1, *p<0.05, **p<0.01, ***p<0.001, ****p<0.0001 for *post hoc* tests.

To determine flexibility of responding following 5 days of self-stimulation training, the lever contingencies were reversed at the beginning of the sixth day such that the formerly active lever became inactive, and *vice versa*. By the fourth day of reversal, both DMS and NAc implantation groups had significantly reversed their lever preference (two-way repeated-measures ANOVA; DMS, significant time x lever interaction, F_(5, 70)_ = 20.06, p <0.0001, no significant effect of time or lever, significant *post hoc* test at day -1, 3, and 5, p <0.05, trending effect at day 4, p <0.1; NAc, F_(5, 70)_ = 5.228, p <0.001, no significant effect of time or lever, trending *post hoc* tests at days -1 and 4; Figure 3A+C). In contrast, mice that self-stimulated DLS dopamine terminals did not significantly alter preference following reversal (no significant effect of lever, time, or time x lever interaction, all p >0.27). When normalized to total number of presses, mice that self-stimulated DMS and NAc dopamine similarly switched their lever preference (two-way repeated-measures ANOVA; DMS, significant time x lever interaction, F_(5, 70)_ = 31.54, p <0.0001, no significant effect of lever or time, significant *post hoc* tests at days -1, 3, and 5, p <0.05, trending effect at day 4, p <0.1; NAc, F_(5, 70)_ = 11.37, p <0.0001, no significant effect of lever or time, significant *post hoc* test at days -1 and 4, p <0.05, trending effect at day 2, p <0.1; Figure 3D+F), while mice that self-stimulated DLS dopamine showed a decrease in initial preference, but no significant preference for the newly active lever (significant time x lever interaction, F_(5, 60)_ = 4.822, p<0.001, trending effect of lever, p <0.1, no significant effect of time, significant *post hoc* test at day -1, p < 0.05, trending effect at day 1, p >0.1; Figure 3E). This was largely replicated in virally-injected DATcre mice: while both DMS and DLS stimulation groups showed a modest tendency to reverse (Supplemental Figure S3A-B), when these values are normalized to % total presses, the DMS group significantly altered preference (Supplemental Figure S3C) while the DLS group did not (Supplemental Figure S3D). Together, these data suggest that dopamine terminal stimulation in DMS and NAc promotes flexible responding for self-stimulation, while DLS terminal stimulation promotes less flexible responding.

**Figure 3.**
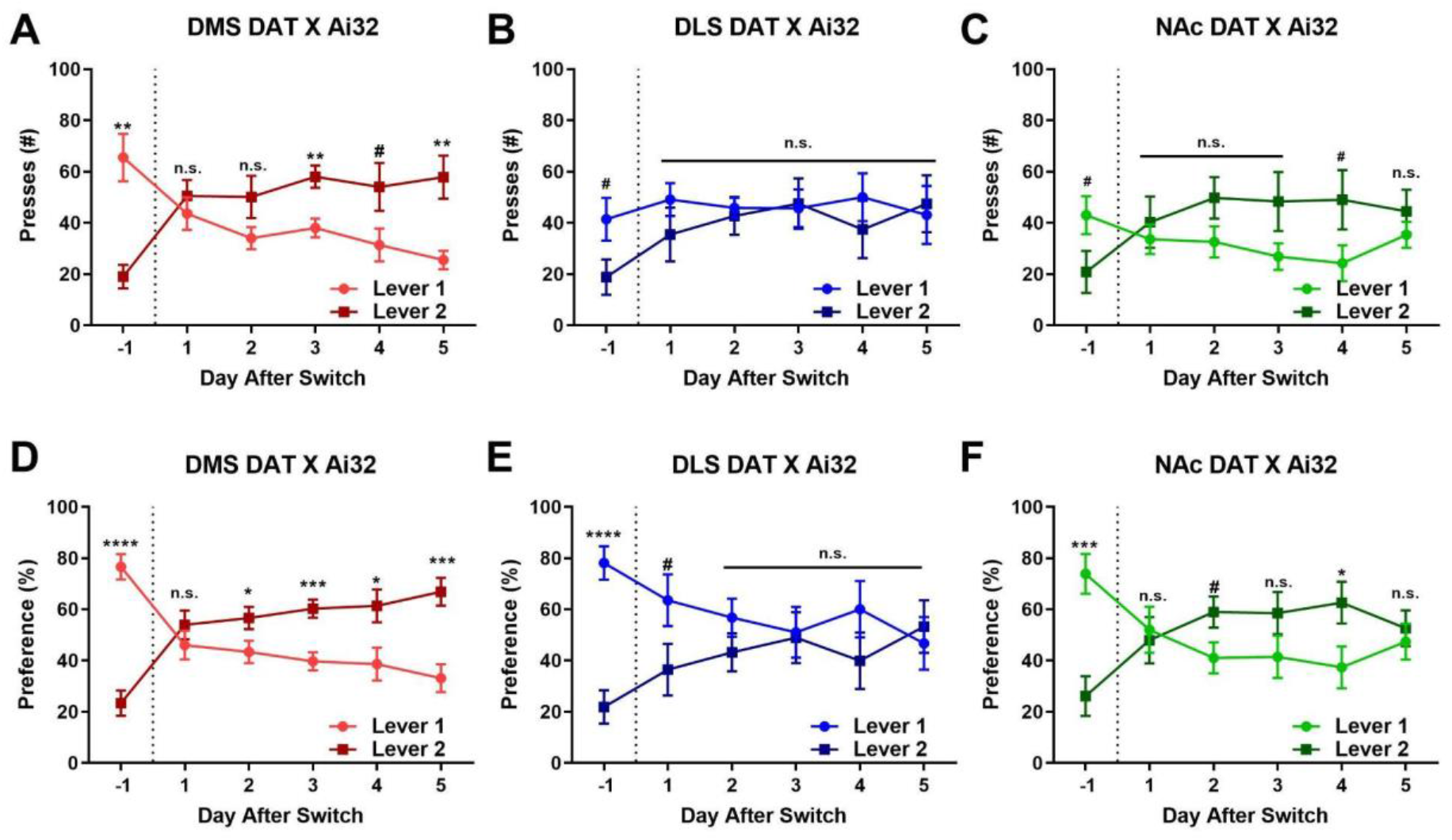
Responding during the reversal phase across implantation sites. (**A-C**) Total number of presses the day before and following contingency reversal (dotted line; lever 1 initially active before becoming inactive and *vice versa*), for mice self-stimulating DMS (**A**), DLS (**B**), and NAc (**C**) dopamine terminals. (**D-F**) Lever preference before and after contingency reversal for mice self-stimulating DMS (**D**), DLS (**E**), and NAc (**F**) dopamine terminals. Mean ± SEM, ^#^p>0.1, *p<0.05, **p<0.01, ***p<0.001, ****p<0.0001 for *post hoc* tests.

### Analysis of Errors and Behavioral Strategy in Reversal

Differences in behavioral flexibility could result from insensitivity to negative outcomes (perseverative errors), or from reverting to a previously-reinforced choice following sampling of the now-correct choice (regressive errors) (Ragozzino, 2007; Floresco et al., 2009; Izquierdo et al., 2017). In order to investigate if error quantity and type differed between groups across training, we quantified the total number of errors within each behavioral session during early (day 1-2) and late (day 3-5) reversal. We found that mice with fiber optics targeting DMS, DLS, or NAc made a partially different number of errors during reversal (two-way repeated-measures ANOVA, trending effect of group; F_(2, 20)_ = 3.288, p= 0.0582, no significant effect of time or group x time interaction; Figure 4A), and this tendency was partially attributed to increased numbers of errors late in reversal for the DLS group relative to the NAc group (trending *post hoc* test DLS vs NAc, p = 0.072, all other tests p >0.1). The vast majority of these errors were attributed to regressive errors, and groups did not differ in numbers of perseverative errors (two-way repeated-measures ANOVA; no significant effect of group, time, or interaction; all p > 0.05; Figure 4B), though groups slightly differed in numbers of regressive errors (two-way repeated-measures ANOVA, trending effect of group; F_(2, 20)_ = 2.774, p = 0.0865; no significant effect of time or time x group interaction) with DLS mice having a weak tendency for higher numbers of regressive errors relative to NAc mice (*post hoc* tests DLS vs NAc, p =0.11 and p =0.12 for early and late, respectively, all other *post hoc* tests p >0.1). Virally-injected DATcre DLS mice similarly made more total, regressive, and perseverative errors, particularly in late reversal, but these results were not statistically significant (Supplemental Figure S4A-C).

**Figure 4.**
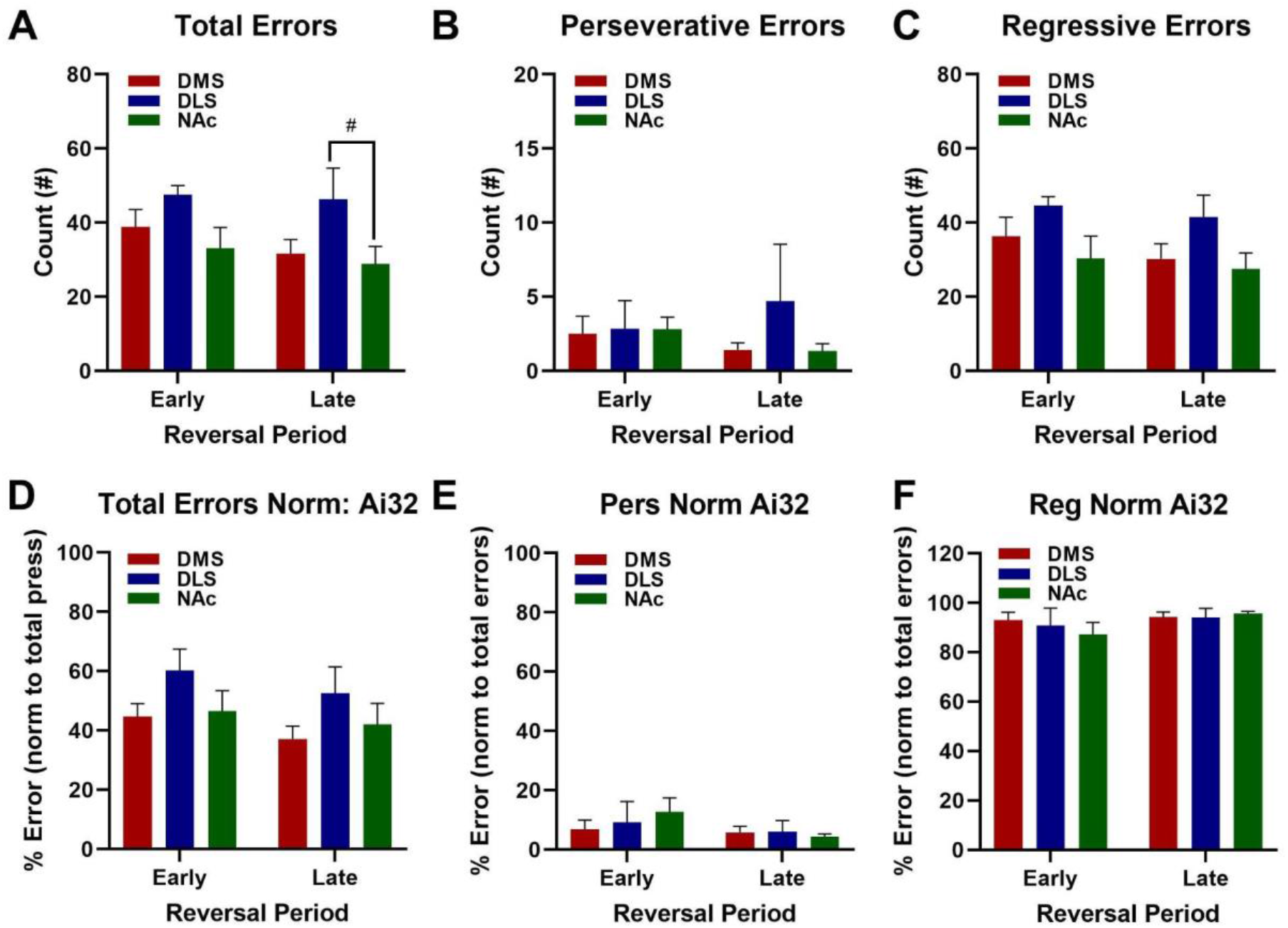
Errors across implantation site groups through early and late reversal. **A+D**. Total number of errors across early (mean of days 1-2) and late (mean of days 3-5) reversal (**A**) and error proportion normalized to total presses (**D**). **B+E**. Number of perseverative errors (errors before sampling of active lever each day) in early and late reversal (**B**) and normalized to total # errors (**E**). **C+F**. Number of regressive errors (errors following sampling of correct lever each day) in early and late reversal (**C**) and normalized to total # errors (**F**). Mean ± SEM # p ≤ 0.1 for *post hoc* tests.

To determine the proportion of errors across variable numbers of responses for each mouse, we next normalized total errors to total numbers of presses, and numbers of perseverative/regressive errors to total numbers of errors. However, groups did not differ in proportion of errors normalized to total presses other than an overall reduction in errors over time (two-way repeated-measures ANOVA; significant effect of time F_(1, 20)_ = 4.436, p <0.05, no significant effect of group or group x time interaction; Figure 4D), and groups did not differ in proportion of perseverative or regressive errors normalized to total error count (two-way repeated-measures ANOVA, no significant effect of group, time, or group x time interaction for perseverative and regressive error proportions; Figure 4E+F). This was also true for virally injected mice, who did not differ in normalized error numbers or error type (two-way repeated-measures ANOVA, no significant effect of group, time, or interaction; Supplemental Figure S4 D-F). Thus, mice self-stimulating DLS dopamine had slightly higher numbers of errors, and these errors tended to be regressive, though this effect was weak.

In addition to error types, elevation of dopamine in different subregions may differently drive reinforcement across learning, which may later influence behavioral strategies following reversal. For example, mice could rely more on their aggregate reinforcement histories experienced during training (win-shift or lose-stay strategies), or on immediate feedback of more proximal outcomes (win-stay or lose-shift strategies; Reed, 2016). Previous work has noted that lesions of DLS reduce lose-shift responding (Skelin et al., 2014), supporting the notion that distinct neural circuits underlie these different behavioral strategies (Gruber & Thapa, 2016). Consistent with this, we found significantly different numbers of lose-stay events following reversal across implantation site groups (two-way repeated-measures ANOVA; significant effect of group; F_(2, 20)_ = 3.99, p < 0.05; no significant effect of time or time x group interaction; Figure 5A), with DLS mice showing significant and partial increases in lose-stay responses late in reversal relative to NAc and DMS mice, respectively (significant *post hoc* test for DLS vs NAc late in reversal, p >0.05, trending *post hoc* test for DLS vs DMS, p =0.059, all other post hoc tests, p <0.1). On the other hand, groups did not differ in numbers of lose-shift, win-stay, or win-shift events (two-way repeated-measures ANOVA, no significant effect of group, all p > 0.05; Figures 5B-D), but mice tended to increase win-stay responding across reversal (partial effect of time, F_(1, 20)_ = 3.770, p <0.1; Figure 5C) and differently altered win-shift responding (partial group x time effect, F_(2, 20)_ = 2.972, p <0.1; Figure 5D; no other significant main effects for lose-stay, win-stay, or win-shift responding, all p <0.1, and no significant *post hoc* tests). Virally-injected DLS mice also had more lose-stay errors, but groups did not differ statistically in numbers of lose-stay, win-stay, win-shift, or lose-shift responses (Supplemental Figure S5A, B+D).

**Figure 5.**
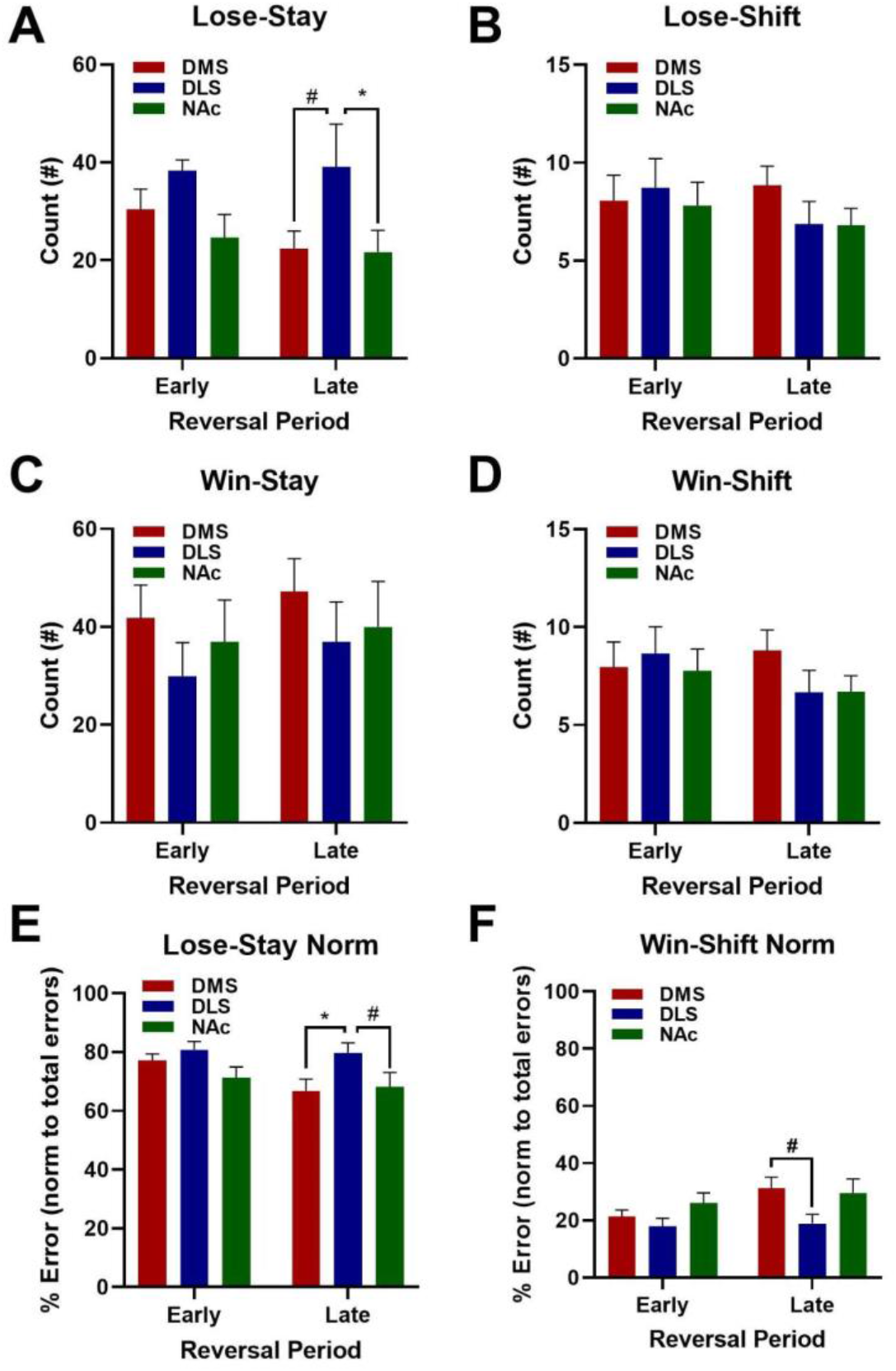
Assessment of behavioral strategy across early and late reversal. **A**. Average number of lose-stay events (selection of inactive lever following selection of inactive lever) following reversal across implantation sites. **B**. Average number of lose-shift events (selection of active lever following selection of inactive lever) following reversal across implantation sites. **C**. Average number of win-stay events (selection of active lever following selection of active lever) following reversal across implantation sites. **D**. Average number of win-shift events (selection of inactive lever following selection of active lever) following reversal across implantation sites. **E**. Average proportion of lose-stay errors normalized to total number of errors. **F**. Average proportion of win-shift errors normalized to total number of errors. Mean ± SEM *, p ≤ 0.05, #, p ≤ 0.1 for *post hoc* tests.

We next normalized error strategies to total number of errors to account for differing number of errors across animals. Implantation site groups had differing proportions of lose-stay errors (two-way repeated-measures ANOVA, significant effect of group, p <0.05, no significant effect of time or group x time interaction; Figure 5E) with DLS having significantly more lose-shift errors than DMS mice (significant *post hoc* for DLS vs DMS, p <0.05), and partially increased lose-shift errors relative to NAc mice (marginally significant *post hoc* test for DLS vs NAc, p <0.1) late in reversal. Groups also slightly differed in their proportion of win-shift errors, which also tended to increase across reversal (two-way repeated-measures ANOVA, trending effect of group and time, F_(2, 20)_ = 3.056 and F_(1, 20)_ = 3.925, respectively, p >0.1, no significant time x group interaction; Figure 5F), which was largely attributed to differences between DMS and DLS groups late in reversal (trending *post hoc* for DMS vs DLS, p =0.054, all other *post hoc* tests p >0.05). Virally-injected mice increased numbers of win-stay responses, normalized lose-stay proportions, and normalized win-shift proportions, suggesting generalized optimization of responding across groups in reversal (Supplemental Figure S5C, E+F), but these effects did not differ between implantation site groups. In sum, this data suggests that self-stimulation of dopamine terminals in DLS resulted in subsequent increases in errors which were largely regressive, increased reliance on lose-stay strategies, and partial decreases in win-shift strategies, which may reflect reliance on initial as opposed to more proximal behavioral outcomes and/or disruption in optimal foraging strategies (Charnov, 1976).

## Discussion

### Distinct roles of dopamine in striatal subregions in behavioral flexibility

The current study suggests that dopamine self-stimulation in dorsomedial (DMS) and accumbens core (NAc) promotes behavioral flexibility, while dorsolateral (DLS) dopamine activation promotes less flexible responding. This finding is consistent with a well-supported model of dorsal striatal functioning, which posits that DMS and DLS promote flexible and inflexible behaviors, respectively (Yin & Knowlton, 2006; Yin et al., 2004; Yin et al., 2009; Thorn et al., 2010; Smith & Graybiel, 2016; Crego et al., 2020); and is consistent with previous dopamine lesion studies suggesting that lesions of DMS or DLS dopamine impair or enhance flexibility, respectively (Faure et al., 2010; Grospe et al., 2018). Dopamine release is initially present in ventromedial striatum early in learning (Willuhn et al., 2012; Radke et al., 2019) before shifting to dorsolateral striatum as behaviors become well-learned (Willuhn et al., 2012). Thus, optogenetic stimulation of DLS dopamine in the current study may bypass normal gating mechanisms of DLS dopamine release to enhance inflexible responding. A similar mechanism may be involved in drug addiction, as drugs of abuse exacerbate habit formation (Nelson & Killcross, 2006; Nelson & Killcross, 2013), potentially by driving release of dopamine directly in DLS (Everitt et al., 2008). Consistent with this notion, cocaine exposure shifts cue-related neuronal activity during discriminative learning from ventral to dorsolateral striatum (Takahashi et al., 2007). Characterizing the brain circuits that guide this progression from ventromedial to dorsolateral dopamine signaling, which may involve reciprocal and spiraling connections between striatum and SNc (Lerner et al., 2015; Ambrosi & Lerner, 2021; Haber et al., 2000), is therefore of great clinical value.

It is possible that optogenetic self-stimulation of DLS dopamine terminals alters responding in reversal via plasticity in DLS (Calabresi et al., 2007), which could promote dominant DLS activity and shift behavior toward inflexible responding (Gremel et al., 2016; O’Hare et al., 2016; Thorn et al., 2010; Yin et al., 2009). DLS receives preferential input from sensorimotor cortical areas implicated in stimulus-response learning, while DMS receives preferential input from associative cortical regions implicated in goal-directed strategies (Suzanne N Haber, 2016). Therefore, dopamine release in DLS may establish dependence on stimulus-response behavior by strengthening cortical-DLS circuits (O’Hare et al., 2016; Gremel et al., 2016). On the other hand, it is possible that activating lateral dopamine enhances generalization between the active and inactive lever, which is supported by slightly lower preference for the active lever prior to reversal in DLS mice (Figure 3B). This is consistent with previous work demonstrating lesions of DLS impair action generalization, while DMS lesions enhance action discrimination (Hilario et al., 2012). One interesting question left unresolved by the current work is how manipulations of dopamine activity in one striatal subregion may influence activity in other striatal subregions once inflexible behaviors have been established. For example, could enhancing DMS/NAc dopamine release following establishment of inflexible behavior restore goal-directed responding? Consistent with this idea, direct deep-brain stimulation of NAc appears to be beneficial in the treatment of drug addiction (Bari et al., 2019) and obsessive-compulsive disorder (Müller et al., 2013).

There are caveats to this model of DMS/DLS functioning: for example, epigenetic manipulations of either DMS or DLS have been sufficient to impair habit formation (Malvaez et al., 2018), which suggests a more general role of dorsal striatum in establishing inflexible responding. Most notably, a recent study found a correlation between DMS (but not DLS) dopamine signaling and compulsive reward-seeking, and demonstrated that optogenetic stimulation of DMS dopamine is sufficient to drive compulsion (Seiler et al., 2020). Similarly, a separate series of studies concluded that self-stimulation of mesolimbic dopamine neurons in ventral tegmental area, which project primarily to ventral striatum, is sufficient to establish compulsive behaviors via strengthening centro-ventral dorsal striatal inputs from orbitofrontal cortex (Pascoli et al., 2015; Pascoli et al., 2018). This may seem to conflict with the current work, however, inflexibility in the current study (continuing to press the inactive lever despite no outcome) differs greatly from compulsivity (persistent behavior despite an aversive outcome). The current work also differs from many other studies of reversal and habit formation due to the use of direct brain stimulation as opposed to natural reinforcers (food or water rewards). Moreover, work from our lab and others have shown that striatal patches - µ-opioid receptor dense portions of striatum that span both DMS and DLS (Jenrette et al., 2019; Nadel et al., 2020) and exhibit unique control over dopamine release (Gerfen, 1992; Graybiel & Ragsdale, 1978) are critical for habit formation. Future work is necessary to investigate how optogenetic stimulation of dopamine within striatal subregions affects habit formation and reversal with natural rewards.

### Optogenetic self-stimulation of distinct striatal subregions alters behavioral strategies

In this study, we noted that mice that self-stimulated dopamine terminals in DLS made more lose-stay and regressive errors following reversal, potentially reflecting a reliance on long-term valuation of the initially reinforced choice. This finding is in agreement with a previous study that found lesioning DMS, and presumably prompting reliance on DLS, resulted in more repetitive responding in a two-choice decision making task (Skelin et al., 2014). On the other hand, Skelin et al. also noted that lesions of DLS result in decreased lose-shift transitions, suggesting DLS is necessary for outcome sensitivity. That both lesions and optogenetic activation of DLS could similarly impair reward sensitivity may seem paradoxical, but it is possible that overactivation of DLS in the current study effectively renders mice outcome-insensitive by driving overreliance on previously learned action valuation. The striatum is thought to encode both object and action-outcome valuation across both dorsal and ventral aspects (Yamada et al., 2013; Kang et al., 2021; Guo et al., 2018; Kim et al., 2015; Yasuda et al., 2012); however, activity in dorsal striatum preferentially encodes valuation of specific actions (Burton et al., 2015; Nakamura et al., 2012; Hollerman et al., 1998), while ventral striatum better tracks outcomes. Studies in primates have also noted long-term value storage in brain centers promoting inflexible responding, including the posterior/tail of the striatum (Kim et al., 2015; Griggs et al., 2017; Guo et al., 2018) and SNr (Yasuda et al., 2012). Additionally, the current study noted a decrease in win-shift errors following self-stimulation of DLS dopamine. Win-shift strategy, and the related phenomenon of spontaneous alternation, both reflect normal and optimal foraging strategies thought to prevent “patch depletion” in foraging animals by varying behavioral choice despite reinforcement (Rayburn-Reeves et al., 2013; Charnov, 1976). Reduced proportion of win-shift events following DLS self-stimulation may reflect breakdown in this normal optimal foraging strategy, which may again reflect overlearning of the initially active lever. Together, these results suggest that enhancing DLS dopamine release may result in long-term activity shifts between the DMS and DLS, resulting in overlearning and overvaluation of the initially reinforced action. Future work could investigate the time course of how DMS and DLS store action valuation, which has not been well characterized.

### The role of dopamine in reversal learning

The current study adds to the body of literature that suggests that dopamine is a key modulator of reversal learning (Izquierdo et al., 2017). Dopamine depletion in Parkinson’s patients (Peterson et al., 2009; Swainson et al., 2000) and in animals treated with 6-OHDA (Grospe et al., 2018; M. O’Neill & Brown, 2007; Clarke et al., 2011) is accompanied by deficits in reversal learning. Moreover, pharmacological modulations of dopamine receptors in NAc (Haluk & Floresco, 2009; Sala-Bayo et al., 2020) and DMS (Sala-Bayo et al., 2020; Wang et al., 2019) impair reversal, and there is a link between D2 receptor availability and reversal learning (Izquierdo et al., 2017). Reversal learning may be facilitated by dopamine signaling positive reward prediction errors (RPE) in ventromedial striatum, which is only noted in animals that are able to successfully reverse (Klanker et al., 2015). Our study is in agreement with this notion, but may also suggest that elevated DLS dopamine during learning later interferes with reversal, which is further supported by a study suggesting DLS inhibition can improve early reversal (Bergstrom et al., 2018).

Previous studies utilizing optogenetic self-stimulation of SNc (Rossi, Sukharnikova, et al., 2013; Ilango et al., 2014) and ventral tegmental dopamine cell bodies (Adamantidis et al., 2011; Ilango et al., 2014) suggest that mice remain capable of rapid reversal following switching of active and inactive levers. This may seem to partially conflict with the current study, however, direct stimulation of cell bodies in midbrain may activate both DMS and DLS projecting dopamine terminals. Consistent with the current work, a study investigating self-stimulation of dopamine terminals in NAc found that mice were flexible following reversal of the active nosepoke (Zell et al., 2020). Future work could explore the difference between stimulating cell bodies and terminals by employing retrograde tracing and intersectional approaches, including Canine adenovirus-2 (Lavoie & Liu, 2020), to perform optogenetics specifically targeting distinct dopamine projection populations (Lerner et al., 2015).

## Conclusion

In summary, the current work supports several previous studies suggesting that selective activation of dopamine is sufficient to reinforce operant behaviors (Adamantidis et al., 2011; Ilango et al., 2014; Keiflin et al., 2019; Covey & Cheer, 2019; Saunders et al., 2018). This work also supports a causative role of DLS dopamine release in establishing inflexible responding. It will be of great clinical significance to extend this work to other forms of flexibility, particularly for natural reinforcers and in drug addiction studies. Finally, future studies should explore how activation of ventromedial dopamine release modifies goal-directed behavior following habit formation, which could yield important clinical insights into therapeutic strategies for mediating disorders characterized by maladaptive habit formation, including obsessive-compulsive disorder and drug addiction.

## Acknowledgements

The authors would like to thank Dr. Tracie Paine for sharing TH antibody and mCherry virus, Dr. Monica Mariani for help with histological techniques, and Nick Hollon for providing insightful feedback on the manuscript. This work was supported by NIH grant 1R15MH122729-0 to CDH. We thank Dr. Adam Eck for fruitful conversations on data analysis with Bonsai. Finally, the authors would also like to thank Lori Lindsay, Forrest Rose, Dorothy Auble, Gigi Knight, Bill Mohler, Mike Miller, Chris Mohler and Laurie Holcomb for research support.

## Supplemental Information

**Supplemental Figure S1.**
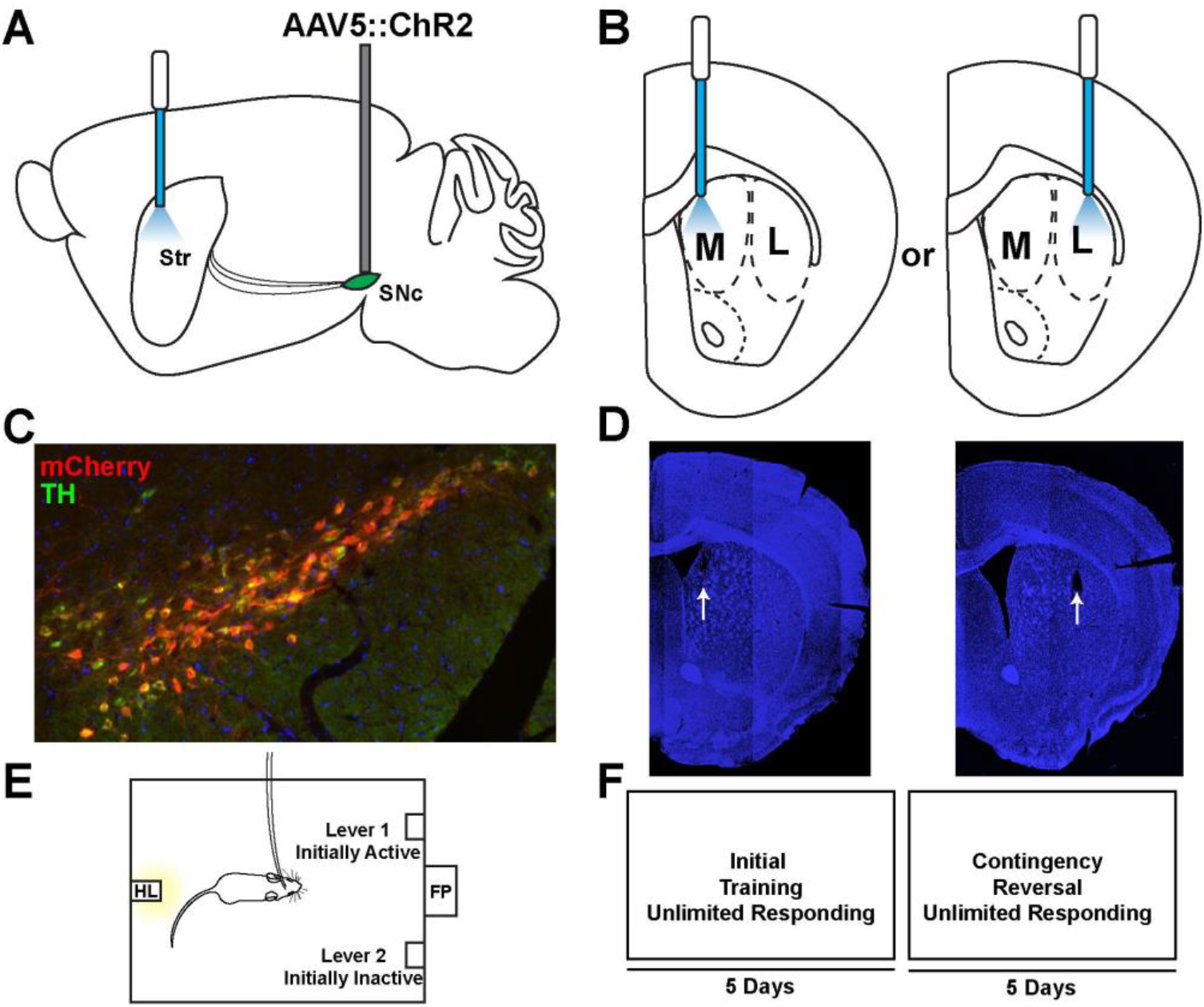
Experimental design for virally-injected DATcre mice. **A-B**. Mice were anesthetized and injected with Cre-dependent AAV:ChR2 before being chronically implanted with fiber optics targeting dorsomedial (**B**, left) or dorsolateral (**B**, right) striatal dopamine terminals. **C**. DATcre mouse midbrain following injection with DIO AAV:ChR2 superimposed over immunohistochemical stain for tyrosine hydroxylase (TH). (**D**) Representative fiber optic implantation sites in DMS (left) and DLS (right). **E**. Operant chamber configuration. Two levers were extended on either side of the food port, and the initially active lever varied across subjects **F**. Schematic of experimental design. Mice could press either lever an unlimited number of times, and following five days of meeting criteria for preferring the active lever (see Methods), the contingency was reversed, and the active lever became inactive and *vice versa* and mice were tested for an additional five days.

**Supplemental Figure S2.**
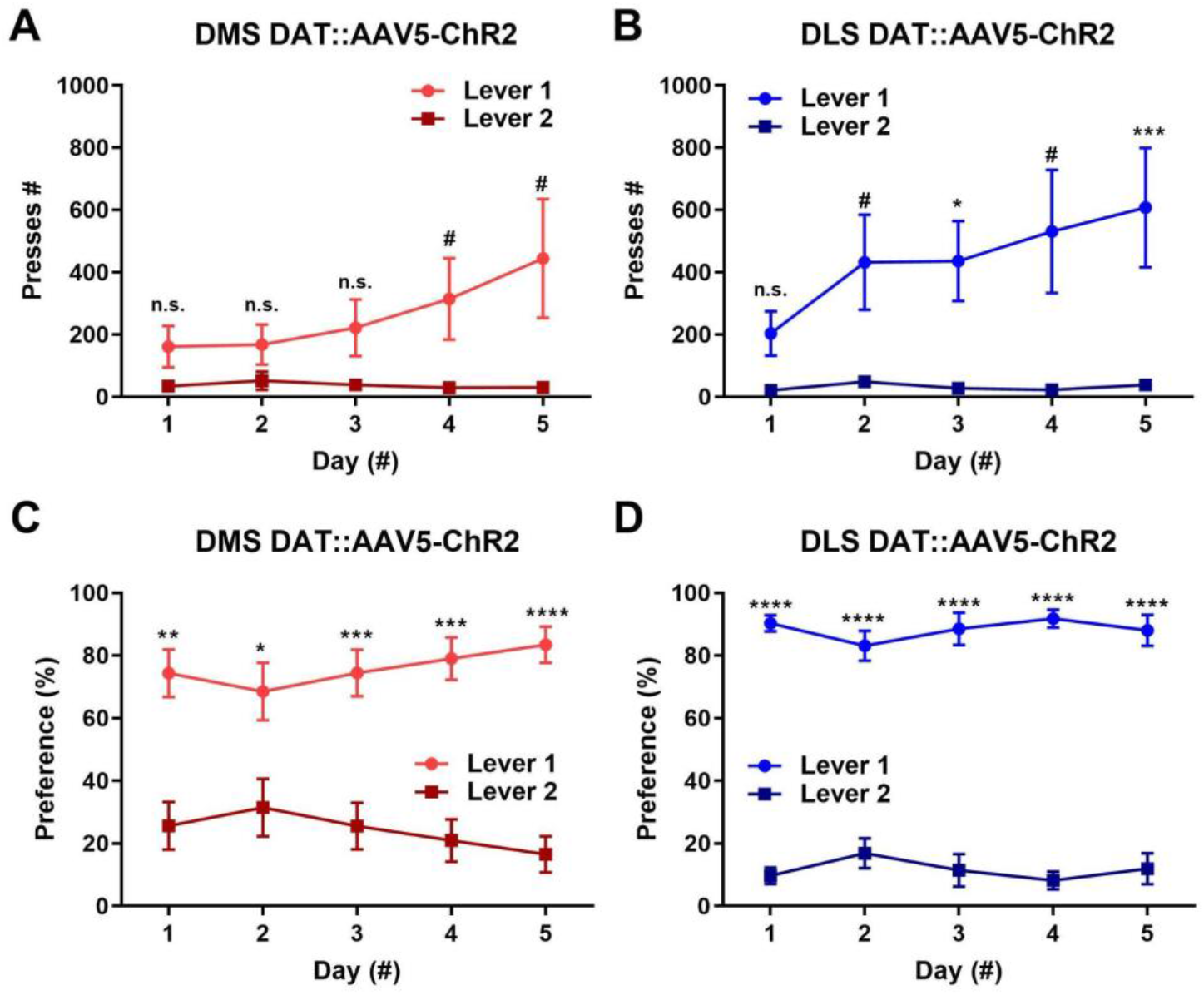
Responding during the initial training phase across implantation sites in virally-injected mice. **A-B**. Total press counts for mice self-stimulating dopamine terminals in DMS (**A;** Two-way repeated measures ANOVA; significant lever x time interaction, F _(4, 40)_ = 3.246, marginal effect of lever, F_(1, 10)_ = 4.78, p <0.01, no significant effect of time, marginally significant *post hoc* tests at days 4-5, p <0.1) and DLS (**B;** significant effect of lever, F_(1, 10)_ = 8.438, p <0.001, no significant lever x time interaction or effect of time, significant *post hoc* tests at days 3+5, p <0.05, trending *post hoc* tests at days 2+4, p <0.1) across the five training days. **C-D**. Press preference (presses on each lever / total presses * 100) for dorsomedial (**C**; two-way repeated measures ANOVA, significant effect of lever, F_(1, 10)_ = 29.37, p <0.001, significant lever x time interaction, F _(4, 40)_ = 5.306, p< 0.01, no significant effect of time, significant *post hoc* tests at all points, p <0.05) and dorsolateral striatum (**D;** significant effect of lever, F_(1, 10)_ = 263.3, p <0.0001, no significant effect of time or time x lever interaction, significant *post hoc* tests at all points, p <0.05) across the five training days. Mean ± SEM. *p<0.05, **p<0.01, ***p<0.001, ****p<0.0001 for *post hoc* tests.

**Supplemental Figure S3.**
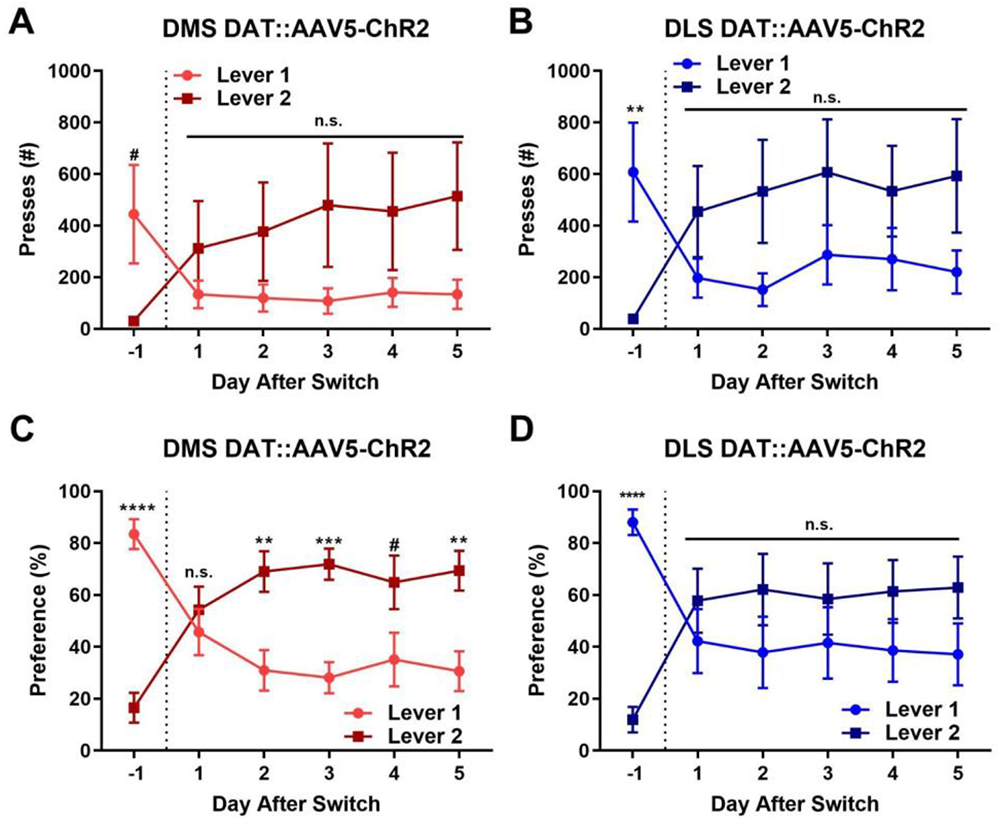
Responding during the reversal phase across implantation sites in virally-injected mice. (**A-B**). Total number of presses the day before and following contingency reversal (dotted line; lever 1 initially active before becoming inactive and *vice versa*), for mice self-stimulating DMS (**A;** Two-way repeated-measures ANOVA, significant time x lever interaction; F_(5, 50)_ = 7.474, p <0.0001, no significant effect of time or lever x time interaction, trending *post hoc* test at day -1, p <0.1) and DLS (**B;** significant time x lever interaction; F_(5, 50)_ = 7.552, p <0.0001, no significant effect of time or lever x time interaction, significant *post hoc* test at day -1, p <0.05) dopamine terminals. (**C-D**). Lever preference before and after contingency reversal for mice self-stimulating DMS (**C;** significant time x lever interaction, F _(5, 50)_ = 22.95, p<0.0001, trending effect of lever, F_(1, 10)_ = 3.800, p <0.1, no significant effect of time, significant *post hoc* tests at day 1, 2, 3, and 5, p <0.05, trending post *hoc test* at day 4, p <0.1) and DLS (**D;** significant time x lever interaction, F _(5, 50)_ = 14.44, p<0.0001, no significant effect of time or lever x time interaction, significant *post hoc* test at day -1, p <0.05) dopamine terminals. Mean ± SEM, *p<0.05, **p<0.01, ***p<0.001, ****p<0.0001 for *post hoc* tests.

**Supplemental Figure S4.**
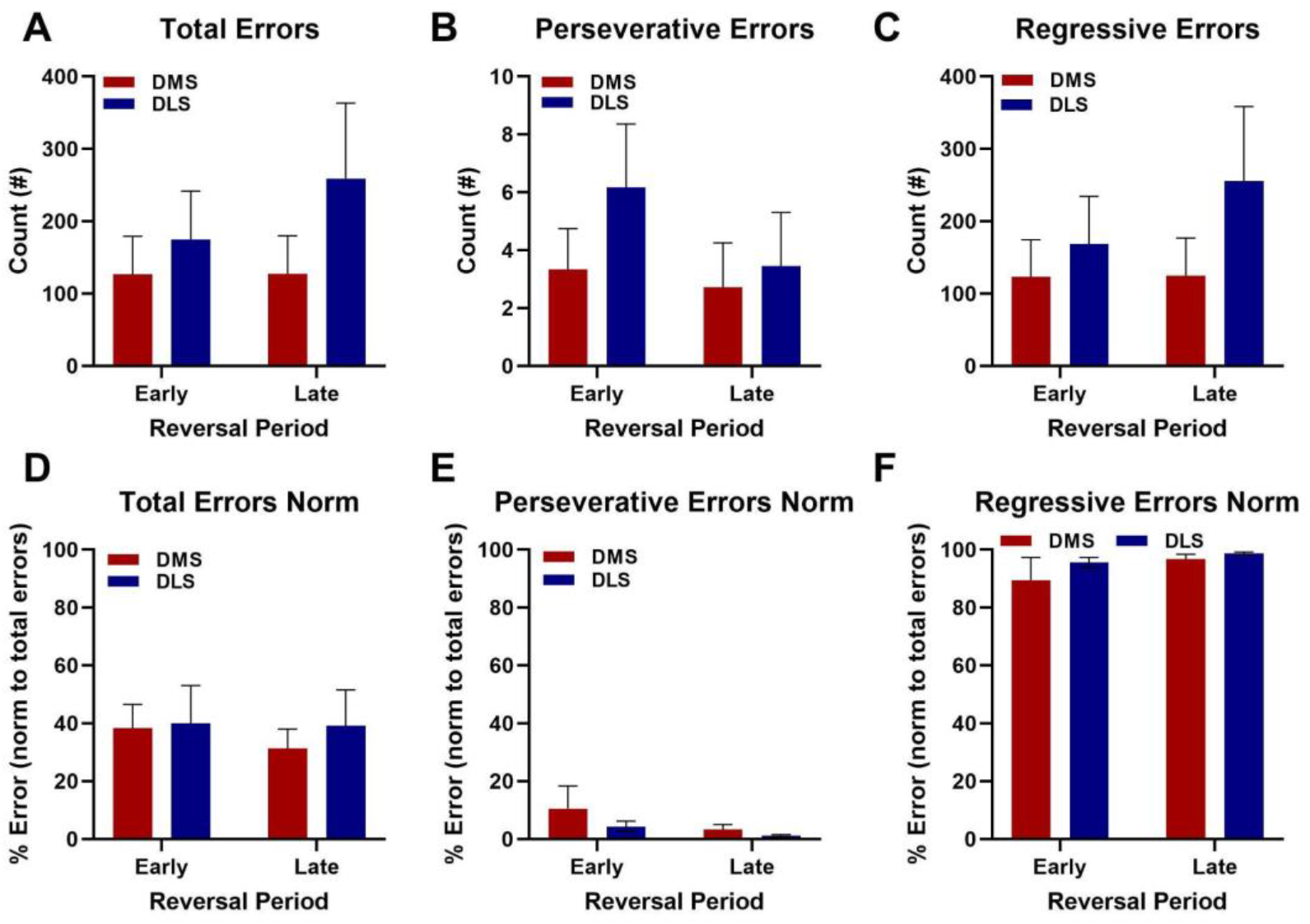
Errors across implantation site groups through early and late reversal in virally-injected mice. **A+D**. Total number of errors (**A**; two-way repeated measures ANOVA; no significant effect of group, time, or group x time interaction; all p>0.05) and errors normalized to total number of presses (**D**; no significant effect of group, time, or group x time interaction; all p>0.05) across DMS and DLS implanted mice. **B+E**. Numbers of perseverative errors (**B, t**wo-way repeated measures ANOVA; no significant effect of group, time, or group x time interaction; all p>0.05) and perseverative errors normalized to total errors (**E**, no significant effect of group, time, or group x time interaction; all p>0.05). **C+F**. Numbers of regressive errors (**C, t**wo-way repeated measures ANOVA; no significant effect of group, time, or group x time interaction; all p>0.05) and regressive errors normalized to total errors across groups (**F**, no significant effect of group, time, or group x time interaction; all p>0.05). Mean ± SEM.

**Supplemental Figure S5.**
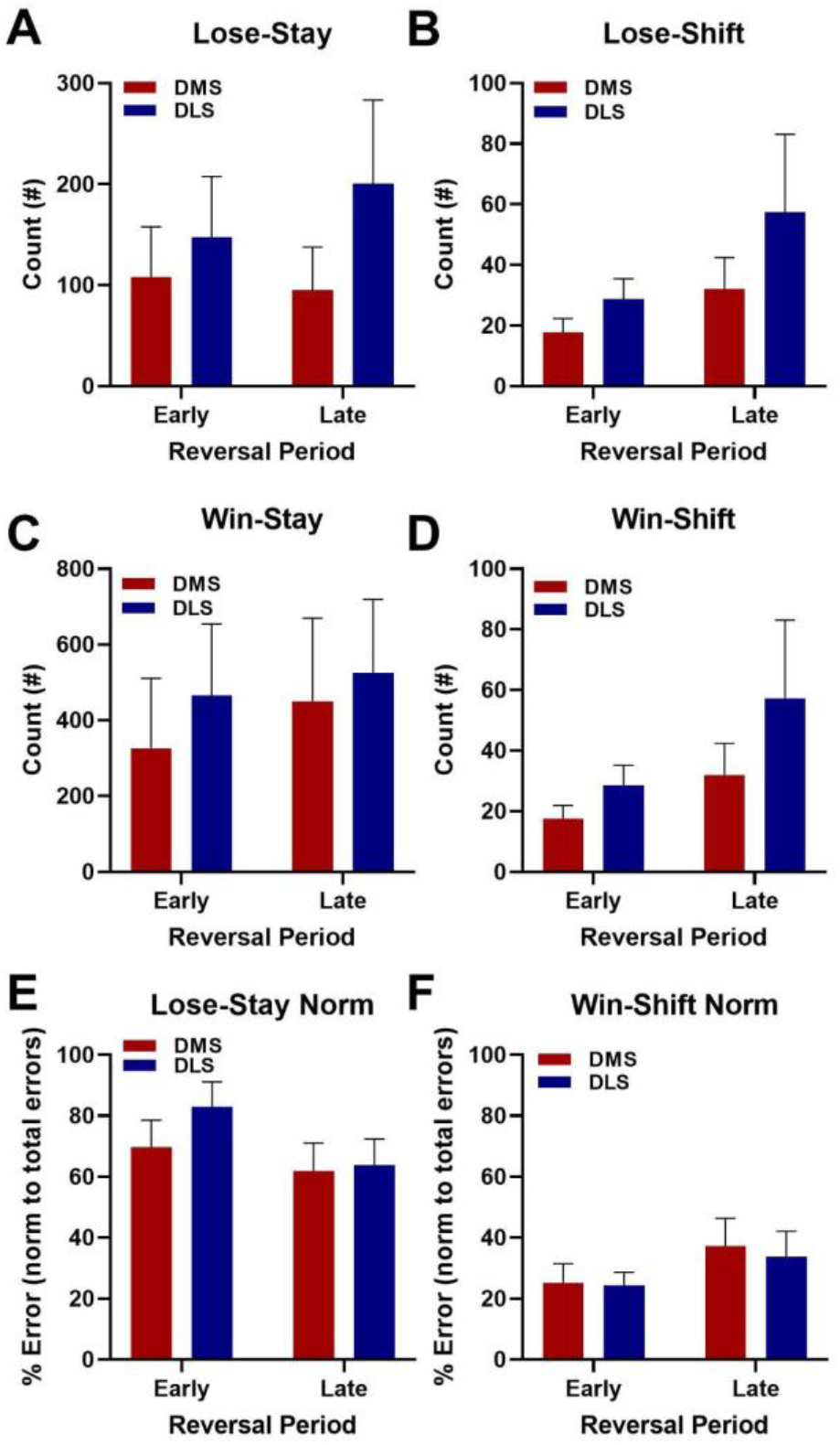
Assessment of behavioral strategy across early and late reversal in virally-injected mice. **A-D**. Number of Lose-Stay (**A**; two-way repeated measures ANOVA; no significant effect of group, time, or interaction; p>0.05), Lose-Shift (**B**; two-way repeated measures ANOVA; no significant effect of group, time, or interaction; p>0.05), Win-Stay (**C**; two-way repeated-measures ANOVA; significant effect of time F_(1, 10)_ = 6.928, p <0.05), and Win-Shift errors (**D**; no significant effect of group, time, or group x time interaction; all p>0.05). **E-F**. Proportion of lose-stay (**E**; two-way repeated-measures ANOVA; significant effect of time F_(1, 10)_ = 5.265, p <0.05, no significant effect of group or group x time interaction) and win-shift errors (**F**; two-way repeated-measures ANOVA; significant effect of time F_(1, 10)_ = 9.302, p <0.05, no significant effect of group or group x time interaction) normalized to total numbers of errors across DMS and DLS implantation sites. Mean ± SEM.

## References

Adamantidis, A. R., Tsai, H.-C., Boutrel, B., Zhang, F., Stuber, G. D., Budygin, E. A., Touriño, C., Bonci, A., Deisseroth, K., & de Lecea, L. (2011). Optogenetic interrogation of dopaminergic modulation of the multiple phases of reward-seeking behavior. The Journal of Neuroscience, 31(30), 10829–10835. https://doi.org/10.1523/JNEUROSCI.2246-11.2011

Adams, C. D., & Dickinson, A. (1981). Instrumental responding following reinforcer devaluation. The Quarterly Journal of Experimental Psychology Section B, 33(2), 109–121. https://doi.org/10.1080/14640748108400816

Ambrosi, P., & Lerner, T. N. (2021). Striatonigrostriatal circuit architecture for disinhibition of dopamine signaling. BioRxiv. https://doi.org/10.1101/2021.06.22.449416

Bäckman, C. M., Malik, N., Zhang, Y., Shan, L., Grinberg, A., Hoffer, B. J., Westphal, H., & Tomac, A. C. (2006). Characterization of a mouse strain expressing Cre recombinase from the 3’ untranslated region of the dopamine transporter locus. Genesis, 44(8), 383–390. https://doi.org/10.1002/dvg.20228

Bari, B. A., Grossman, C. D., Lubin, E. E., Rajagopalan, A. E., Cressy, J. I., & Cohen, J. Y. (2019). Stable representations of decision variables for flexible behavior. Neuron, 103(5), 922-933.e7. https://doi.org/10.1016/j.neuron.2019.06.001

Bergstrom, H. C., Lipkin, A. M., Lieberman, A. G., Pinard, C. R., Gunduz-Cinar, O., Brockway, E. T., Taylor, W. W., Nonaka, M., Bukalo, O., Wills, T. A., Rubio, F. J., Li, X., Pickens, C. L., Winder, D. G., & Holmes, A. (2018). Dorsolateral Striatum Engagement Interferes with Early Discrimination Learning. Cell Reports, 23(8), 2264–2272. https://doi.org/10.1016/j.celrep.2018.04.081

Burton, A. C., Nakamura, K., & Roesch, M. R. (2015). From ventral-medial to dorsal-lateral striatum: neural correlates of reward-guided decision-making. Neurobiology of Learning and Memory, 117, 51–59. https://doi.org/10.1016/j.nlm.2014.05.003

Calabresi, P., Picconi, B., Tozzi, A., & Di Filippo, M. (2007). Dopamine-mediated regulation of corticostriatal synaptic plasticity. Trends in Neurosciences, 30(5), 211–219. https://doi.org/10.1016/j.tins.2007.03.001

Castañé, A., Theobald, D. E. H., & Robbins, T. W. (2010). Selective lesions of the dorsomedial striatum impair serial spatial reversal learning in rats. Behavioural Brain Research, 210(1), 74–83. https://doi.org/10.1016/j.bbr.2010.02.017

Charnov, E. L. (1976). Optimal foraging, the marginal value theorem. Theoretical Population Biology, 9(2), 129–136. https://doi.org/10.1016/0040-5809(76)90040-X

Clarke, H. F., Hill, G. J., Robbins, T. W., & Roberts, A. C. (2011). Dopamine, but not serotonin, regulates reversal learning in the marmoset caudate nucleus. The Journal of Neuroscience, 31(11), 4290–4297. https://doi.org/10.1523/JNEUROSCI.5066-10.2011

Clark, L., Cools, R., & Robbins, T. W. (2004). The neuropsychology of ventral prefrontal cortex: decision-making and reversal learning. Brain and Cognition, 55(1), 41–53. https://doi.org/10.1016/S0278-2626(03)00284-7

Covey, D. P., & Cheer, J. F. (2019). Accumbal Dopamine Release Tracks the Expectation of Dopamine Neuron-Mediated Reinforcement. Cell Reports, 27(2), 481-490.e3. https://doi.org/10.1016/j.celrep.2019.03.055

Crego, A. C. G., Štoček, F., Marchuk, A. G., Carmichael, J. E., van der Meer, M. A. A., & Smith, K. S. (2020). Complementary Control over Habits and Behavioral Vigor by Phasic Activity in the Dorsolateral Striatum. The Journal of Neuroscience, 40(10), 2139–2153. https://doi.org/10.1523/JNEUROSCI.1313-19.2019

Everitt, B. J., Belin, D., Economidou, D., Pelloux, Y., Dalley, J. W., & Robbins, T. W. (2008). Review. Neural mechanisms underlying the vulnerability to develop compulsive drug-seeking habits and addiction. Philosophical Transactions of the Royal Society of London. Series B, Biological Sciences, 363(1507), 3125–3135. https://doi.org/10.1098/rstb.2008.0089

Faure, A, Leblanc-Veyrac, P., & El Massioui, N. (2010). Dopamine agonists increase perseverative instrumental responses but do not restore habit formation in a rat model of Parkinsonism. Neuroscience, 168(2), 477–486. https://doi.org/10.1016/j.neuroscience.2010.03.047

Faure, Alexis, Haberland, U., Condé, F., & El Massioui, N. (2005). Lesion to the nigrostriatal dopamine system disrupts stimulus-response habit formation. The Journal of Neuroscience, 25(11), 2771–2780. https://doi.org/10.1523/JNEUROSCI.3894-04.2005

Floresco, S. B., Ghods-Sharifi, S., Vexelman, C., & Magyar, O. (2006). Dissociable roles for the nucleus accumbens core and shell in regulating set shifting. The Journal of Neuroscience, 26(9), 2449–2457. https://doi.org/10.1523/JNEUROSCI.4431-05.2006

Floresco, S. B., Zhang, Y., & Enomoto, T. (2009). Neural circuits subserving behavioral flexibility and their relevance to schizophrenia. Behavioural Brain Research, 204(2), 396–409. https://doi.org/10.1016/j.bbr.2008.12.001

Gerfen, C. R. (1992). The neostriatal mosaic: multiple levels of compartmental organization. Journal of Neural Transmission. Supplementum, 36, 43–59. https://doi.org/10.1007/978-3-7091-9211-5_4

Gillan, C. M., Papmeyer, M., Morein-Zamir, S., Sahakian, B. J., Fineberg, N. A., Robbins, T. W., & de Wit, S. (2011). Disruption in the balance between goal-directed behavior and habit learning in obsessive-compulsive disorder. The American Journal of Psychiatry, 168(7), 718–726. https://doi.org/10.1176/appi.ajp.2011.10071062

Graybiel, A. M., & Ragsdale, C. W. (1978). Histochemically distinct compartments in the striatum of human, monkeys, and cat demonstrated by acetylthiocholinesterase staining. Proceedings of the National Academy of Sciences of the United States of America, 75(11), 5723–5726. https://doi.org/10.1073/pnas.75.11.5723

Gremel, C. M., Chancey, J. H., Atwood, B. K., Luo, G., Neve, R., Ramakrishnan, C., Deisseroth, K., Lovinger, D. M., & Costa, R. M. (2016). Endocannabinoid modulation of orbitostriatal circuits gates habit formation. Neuron, 90(6), 1312–1324. https://doi.org/10.1016/j.neuron.2016.04.043

Gremel, C. M., & Costa, R. M. (2013). Orbitofrontal and striatal circuits dynamically encode the shift between goal-directed and habitual actions. Nature Communications, 4, 2264. https://doi.org/10.1038/ncomms3264

Griggs, W. S., Kim, H. F., Ghazizadeh, A., Costello, M. G., Wall, K. M., & Hikosaka, O. (2017). Flexible and stable value coding areas in caudate head and tail receive anatomically distinct cortical and subcortical inputs. Frontiers in Neuroanatomy, 11, 106. https://doi.org/10.3389/fnana.2017.00106

Groman, S. M., Lee, B., London, E. D., Mandelkern, M. A., James, A. S., Feiler, K., Rivera, R., Dahlbom, M., Sossi, V., Vandervoort, E., & Jentsch, J. D. (2011). Dorsal striatal D2-like receptor availability covaries with sensitivity to positive reinforcement during discrimination learning. The Journal of Neuroscience, 31(20), 7291–7299. https://doi.org/10.1523/JNEUROSCI.0363-11.2011

Grospe, G. M., Baker, P. M., & Ragozzino, M. E. (2018). Cognitive Flexibility Deficits Following 6-OHDA Lesions of the Rat Dorsomedial Striatum. Neuroscience, 374, 80–90. https://doi.org/10.1016/j.neuroscience.2018.01.032

Gruber, A. J., & Thapa, R. (2016). The Memory Trace Supporting Lose-Shift Responding Decays Rapidly after Reward Omission and Is Distinct from Other Learning Mechanisms in Rats. ENeuro, 3(6). https://doi.org/10.1523/ENEURO.0167-16.2016

Guo, L., Walker, W. I., Ponvert, N. D., Penix, P. L., & Jaramillo, S. (2018). Stable representation of sounds in the posterior striatum during flexible auditory decisions. Nature Communications, 9(1), 1534. https://doi.org/10.1038/s41467-018-03994-3

Haber, Suzanne N. (2016). Corticostriatal circuitry. Dialogues in Clinical Neuroscience, 18(1), 7–21. https://doi.org/10.1007/978-1-4939-3474-4_135

Haber, S N, Fudge, J. L., & McFarland, N. R. (2000). Striatonigrostriatal pathways in primates form an ascending spiral from the shell to the dorsolateral striatum. The Journal of Neuroscience, 20(6), 2369–2382. https://doi.org/10.1523/JNEUROSCI.20-06-02369.2000

Haluk, D. M., & Floresco, S. B. (2009). Ventral striatal dopamine modulation of different forms of behavioral flexibility. Neuropsychopharmacology, 34(8), 2041–2052. https://doi.org/10.1038/npp.2009.21

Hilario, M., Holloway, T., Jin, X., & Costa, R. M. (2012). Different dorsal striatum circuits mediate action discrimination and action generalization. The European Journal of Neuroscience, 35(7), 1105–1114. https://doi.org/10.1111/j.1460-9568.2012.08073.x

Hollerman, J. R., Tremblay, L., & Schultz, W. (1998). Influence of reward expectation on behavior-related neuronal activity in primate striatum. Journal of Neurophysiology, 80(2), 947–963. https://doi.org/10.1152/jn.1998.80.2.947

Howard, C. D., Li, H., Geddes, C. E., & Jin, X. (2017). Dynamic nigrostriatal dopamine biases action selection. Neuron, 93(6), 1436-1450.e8. https://doi.org/10.1016/j.neuron.2017.02.029

Ilango, A., Kesner, A. J., Keller, K. L., Stuber, G. D., Bonci, A., & Ikemoto, S. (2014). Similar roles of substantia nigra and ventral tegmental dopamine neurons in reward and aversion. The Journal of Neuroscience, 34(3), 817–822. https://doi.org/10.1523/JNEUROSCI.1703-13.2014

Izquierdo, A., Brigman, J. L., Radke, A. K., Rudebeck, P. H., & Holmes, A. (2017). The neural basis of reversal learning: An updated perspective. Neuroscience, 345, 12–26. https://doi.org/10.1016/j.neuroscience.2016.03.021

Jenrette, T. A., Logue, J. B., & Horner, K. A. (2019). Lesions of the Patch Compartment of Dorsolateral Striatum Disrupt Stimulus-Response Learning. Neuroscience, 415, 161–172. https://doi.org/10.1016/j.neuroscience.2019.07.033

Kang, J., Kim, H., Hwang, S. H., Han, M., Lee, S.-H., & Kim, H. F. (2021). Primate ventral striatum maintains neural representations of the value of previously rewarded objects for habitual seeking. Nature Communications, 12(1), 2100. https://doi.org/10.1038/s41467-021-22335-5

Keiflin, R., Pribut, H. J., Shah, N. B., & Janak, P. H. (2019). Ventral tegmental dopamine neurons participate in reward identity predictions. Current Biology, 29(1), 93-103.e3. https://doi.org/10.1016/j.cub.2018.11.050

Kim, H. F., Ghazizadeh, A., & Hikosaka, O. (2015). Dopamine Neurons Encoding Long-Term Memory of Object Value for Habitual Behavior. Cell, 163(5), 1165–1175. https://doi.org/10.1016/j.cell.2015.10.063

Klanker, M., Sandberg, T., Joosten, R., Willuhn, I., Feenstra, M., & Denys, D. (2015). Phasic dopamine release induced by positive feedback predicts individual differences in reversal learning. Neurobiology of Learning and Memory, 125, 135–145. https://doi.org/10.1016/j.nlm.2015.08.011

Lavoie, A., & Liu, B.-H. (2020). Canine adenovirus 2: A natural choice for brain circuit dissection. Frontiers in Molecular Neuroscience, 13, 9. https://doi.org/10.3389/fnmol.2020.00009

Leeson, V. C., Robbins, T. W., Matheson, E., Hutton, S. B., Ron, M. A., Barnes, T. R. E., & Joyce, E. M. (2009). Discrimination learning, reversal, and set-shifting in first-episode schizophrenia: stability over six years and specific associations with medication type and disorganization syndrome. Biological Psychiatry, 66(6), 586–593. https://doi.org/10.1016/j.biopsych.2009.05.016

Lerner, T. N., Shilyansky, C., Davidson, T. J., Evans, K. E., Beier, K. T., Zalocusky, K. A., Crow, A. K., Malenka, R. C., Luo, L., Tomer, R., & Deisseroth, K. (2015). Intact-Brain Analyses Reveal Distinct Information Carried by SNc Dopamine Subcircuits. Cell, 162(3), 635–647. https://doi.org/10.1016/j.cell.2015.07.014

Madisen, L., Mao, T., Koch, H., Zhuo, J., Berenyi, A., Fujisawa, S., Hsu, Y.-W. A., Garcia, A. J., Gu, X., Zanella, S., Kidney, J., Gu, H., Mao, Y., Hooks, B. M., Boyden, E. S., Buzsáki, G., Ramirez, J. M., Jones, A. R., Svoboda, K., … Zeng, H. (2012). A toolbox of Cre-dependent optogenetic transgenic mice for light-induced activation and silencing. Nature Neuroscience, 15(5), 793–802. https://doi.org/10.1038/nn.3078

Malvaez, M., Greenfield, V. Y., Matheos, D. P., Angelillis, N. A., Murphy, M. D., Kennedy, P. J., Wood, M. A., & Wassum, K. M. (2018). Habits are negatively regulated by histone deacetylase 3 in the dorsal striatum. Biological Psychiatry, 84(5), 383–392. https://doi.org/10.1016/j.biopsych.2018.01.025

Mink, J. W. (2003). The Basal Ganglia and involuntary movements: impaired inhibition of competing motor patterns. Archives of Neurology, 60(10), 1365–1368. https://doi.org/10.1001/archneur.60.10.1365

Müller, U. J., Voges, J., Steiner, J., Galazky, I., Heinze, H.-J., Möller, M., Pisapia, J., Halpern, C., Caplan, A., Bogerts, B., & Kuhn, J. (2013). Deep brain stimulation of the nucleus accumbens for the treatment of addiction. Annals of the New York Academy of Sciences, 1282, 119–128. https://doi.org/10.1111/j.1749-6632.2012.06834.x

Nadel, J. A., Pawelko, S. S., Copes-Finke, D., Neidhart, M., & Howard, C. D. (2020). Lesion of striatal patches disrupts habitual behaviors and increases behavioral variability. Plos One, 15(1), e0224715. https://doi.org/10.1371/journal.pone.0224715

Nakamura, K., Santos, G. S., Matsuzaki, R., & Nakahara, H. (2012). Differential reward coding in the subdivisions of the primate caudate during an oculomotor task. The Journal of Neuroscience, 32(45), 15963–15982. https://doi.org/10.1523/JNEUROSCI.1518-12.2012

Nelson, A., & Killcross, S. (2006). Amphetamine exposure enhances habit formation. The Journal of Neuroscience, 26(14), 3805–3812. https://doi.org/10.1523/JNEUROSCI.4305-05.2006

Nelson, A. J. D., & Killcross, S. (2013). Accelerated habit formation following amphetamine exposure is reversed by D1, but enhanced by D2, receptor antagonists. Frontiers in Neuroscience, 7, 76. https://doi.org/10.3389/fnins.2013.00076

O’Hare, J. K., Ade, K. K., Sukharnikova, T., Van Hooser, S. D., Palmeri, M. L., Yin, H. H., & Calakos, N. (2016). Pathway-Specific Striatal Substrates for Habitual Behavior. Neuron, 89(3), 472–479. https://doi.org/10.1016/j.neuron.2015.12.032

O’Neill, B., Patel, J. C., & Rice, M. E. (2017). Characterization of Optically and Electrically Evoked Dopamine Release in Striatal Slices from Digenic Knock-in Mice with DAT-Driven Expression of Channelrhodopsin. ACS Chemical Neuroscience, 8(2), 310–319. https://doi.org/10.1021/acschemneuro.6b00300

O’Neill, M., & Brown, V. J. (2007). The effect of striatal dopamine depletion and the adenosine A2A antagonist KW-6002 on reversal learning in rats. Neurobiology of Learning and Memory, 88(1), 75–81. https://doi.org/10.1016/j.nlm.2007.03.003

Pascoli, V., Hiver, A., Van Zessen, R., Loureiro, M., Achargui, R., Harada, M., Flakowski, J., & Lüscher, C. (2018). Stochastic synaptic plasticity underlying compulsion in a model of addiction. Nature, 564(7736), 366–371. https://doi.org/10.1038/s41586-018-0789-4

Pascoli, V., Terrier, J., Hiver, A., & Lüscher, C. (2015). Sufficiency of mesolimbic dopamine neuron stimulation for the progression to addiction. Neuron, 88(5), 1054–1066. https://doi.org/10.1016/j.neuron.2015.10.017

Peterson, D. A., Elliott, C., Song, D. D., Makeig, S., Sejnowski, T. J., & Poizner, H. (2009). Probabilistic reversal learning is impaired in Parkinson’s disease. Neuroscience, 163(4), 1092–1101. https://doi.org/10.1016/j.neuroscience.2009.07.033

Radke, A. K., Kocharian, A., Covey, D. P., Lovinger, D. M., Cheer, J. F., Mateo, Y., & Holmes, A. (2019). Contributions of nucleus accumbens dopamine to cognitive flexibility. The European Journal of Neuroscience, 50(3), 2023–2035. https://doi.org/10.1111/ejn.14152

Ragozzino, M. E. (2007). The contribution of the medial prefrontal cortex, orbitofrontal cortex, and dorsomedial striatum to behavioral flexibility. Annals of the New York Academy of Sciences, 1121, 355–375. https://doi.org/10.1196/annals.1401.013

Rayburn-Reeves, R. M., Stagner, J. P., Kirk, C. R., & Zentall, T. R. (2013). Reversal learning in rats (Rattus norvegicus) and pigeons (Columba livia): qualitative differences in behavioral flexibility. Journal of Comparative Psychology, 127(2), 202–211. https://doi.org/10.1037/a0026311

Reed, P. (2016). Win-stay and win-shift lever-press strategies in an appetitively reinforced task for rats. Learning & Behavior, 44(4), 340–346. https://doi.org/10.3758/s13420-016-0225-2

Rossi, M. A., Fan, D., Barter, J. W., & Yin, H. H. (2013). Bidirectional modulation of substantia nigra activity by motivational state. Plos One, 8(8), e71598. https://doi.org/10.1371/journal.pone.0071598

Rossi, M. A., Sukharnikova, T., Hayrapetyan, V. Y., Yang, L., & Yin, H. H. (2013). Operant self-stimulation of dopamine neurons in the substantia nigra. Plos One, 8(6), e65799. https://doi.org/10.1371/journal.pone.0065799

Sala-Bayo, J., Fiddian, L., Nilsson, S. R. O., Hervig, M. E., McKenzie, C., Mareschi, A., Boulos, M., Zhukovsky, P., Nicholson, J., Dalley, J. W., Alsiö, J., & Robbins, T. W. (2020). Dorsal and ventral striatal dopamine D1 and D2 receptors differentially modulate distinct phases of serial visual reversal learning. Neuropsychopharmacology, 45(5), 736–744. https://doi.org/10.1038/s41386-020-0612-4

Saunders, B. T., Richard, J. M., Margolis, E. B., & Janak, P. H. (2018). Dopamine neurons create Pavlovian conditioned stimuli with circuit-defined motivational properties. Nature Neuroscience, 21(8), 1072–1083. https://doi.org/10.1038/s41593-018-0191-4

Schultz, W. (1998). Predictive reward signal of dopamine neurons. Journal of Neurophysiology, 80(1), 1–27. https://doi.org/10.1152/jn.1998.80.1.1

Seiler, J. L., Cosme, C. V., Sherathiya, V. N., Bianco, J. M., & Lerner, T. N. (2020). Dopamine signaling in the dorsomedial striatum promotes compulsive behavior. BioRxiv. https://doi.org/10.1101/2020.03.30.016238

Skelin, I., Hakstol, R., VanOyen, J., Mudiayi, D., Molina, L. A., Holec, V., Hong, N. S., Euston, D. R., McDonald, R. J., & Gruber, A. J. (2014). Lesions of dorsal striatum eliminate lose-switch responding but not mixed-response strategies in rats. The European Journal of Neuroscience, 39(10), 1655–1663. https://doi.org/10.1111/ejn.12518

Smith, K. S., & Graybiel, A. M. (2016). Habit formation coincides with shifts in reinforcement representations in the sensorimotor striatum. Journal of Neurophysiology, 115(3), 1487–1498. https://doi.org/10.1152/jn.00925.2015

Stern, C. E., & Passingham, R. E. (1995). The nucleus accumbens in monkeys (Macaca fascicularis). III. Reversal learning. Experimental Brain Research, 106(2), 239–247. https://doi.org/10.1007/BF00241119

Swainson, R., Rogers, R. D., Sahakian, B. J., Summers, B. A., Polkey, C. E., & Robbins, T. W. (2000). Probabilistic learning and reversal deficits in patients with Parkinson’s disease or frontal or temporal lobe lesions: possible adverse effects of dopaminergic medication. Neuropsychologia, 38(5), 596–612. https://doi.org/10.1016/S0028-3932(99)00103-7

Takahashi, Y., Roesch, M. R., Stalnaker, T. A., & Schoenbaum, G. (2007). Cocaine exposure shifts the balance of associative encoding from ventral to dorsolateral striatum. Frontiers in Integrative Neuroscience, 1(11). https://doi.org/10.3389/neuro.07/011.2007

Thorn, C. A., Atallah, H., Howe, M., & Graybiel, A. M. (2010). Differential dynamics of activity changes in dorsolateral and dorsomedial striatal loops during learning. Neuron, 66(5), 781–795. https://doi.org/10.1016/j.neuron.2010.04.036

Tsai, H.-C., Zhang, F., Adamantidis, A., Stuber, G. D., Bonci, A., de Lecea, L., & Deisseroth, K. (2009). Phasic firing in dopaminergic neurons is sufficient for behavioral conditioning. Science, 324(5930), 1080–1084. https://doi.org/10.1126/science.1168878

Wang, X., Qiao, Y., Dai, Z., Sui, N., Shen, F., Zhang, J., & Liang, J. (2019). Medium spiny neurons of the anterior dorsomedial striatum mediate reversal learning in a cell-type-dependent manner. Brain Structure & Function, 224(1), 419–434. https://doi.org/10.1007/s00429-018-1780-4

Willuhn, I., Burgeno, L. M., Everitt, B. J., & Phillips, P. E. M. (2012). Hierarchical recruitment of phasic dopamine signaling in the striatum during the progression of cocaine use. Proceedings of the National Academy of Sciences of the United States of America, 109(50), 20703–20708. https://doi.org/10.1073/pnas.1213460109

Yamada, H., Inokawa, H., Matsumoto, N., Ueda, Y., Enomoto, K., & Kimura, M. (2013). Coding of the long-term value of multiple future rewards in the primate striatum. Journal of Neurophysiology, 109(4), 1140–1151. https://doi.org/10.1152/jn.00289.2012

Yasuda, M., Yamamoto, S., & Hikosaka, O. (2012). Robust representation of stable object values in the oculomotor Basal Ganglia. The Journal of Neuroscience, 32(47), 16917–16932. https://doi.org/10.1523/JNEUROSCI.3438-12.2012

Yin, H. H., Knowlton, B. J., & Balleine, B. W. (2004). Lesions of dorsolateral striatum preserve outcome expectancy but disrupt habit formation in instrumental learning. The European Journal of Neuroscience, 19(1), 181–189. https://doi.org/10.1111/j.1460-9568.2004.03095.x

Yin, H. H., & Knowlton, B. J. (2006). The role of the basal ganglia in habit formation. Nature Reviews. Neuroscience, 7(6), 464–476. https://doi.org/10.1038/nrn1919

Yin, H. H., Mulcare, S. P., Hilário, M. R. F., Clouse, E., Holloway, T., Davis, M. I., Hansson, A. C., Lovinger, D. M., & Costa, R. M. (2009). Dynamic reorganization of striatal circuits during the acquisition and consolidation of a skill. Nature Neuroscience, 12(3), 333–341. https://doi.org/10.1038/nn.2261

Yin, H. H., Ostlund, S. B., Knowlton, B. J., & Balleine, B. W. (2005). The role of the dorsomedial striatum in instrumental conditioning. The European Journal of Neuroscience, 22(2), 513–523. https://doi.org/10.1111/j.1460-9568.2005.04218.x

Zell, V., Steinkellner, T., Hollon, N. G., Warlow, S. M., Souter, E., Faget, L., Hunker, A. C., Jin, X., Zweifel, L. S., & Hnasko, T. S. (2020). VTA Glutamate Neuron Activity Drives Positive Reinforcement Absent Dopamine Co-release. Neuron, 107(5), 864-873.e4. https://doi.org/10.1016/j.neuron.2020.06.011

